## Supplemental file_Iglesias Olle for "The *Wnt/β-catenin/TCF/Sp5/Zic4* gene network that regulates head organizer activity in *Hydra* is differentially regulated in epidermis and gastrodermis": Iglesias-Olle_Sp5-regulation_SUPPLEMENT_20240422.pdf

### SUPPLEMENTARY TABLES

**TABLE S1:** Accession numbers (AN) of the *Hydra* genes used in this study p2

**TABLE S2:** List of primers, siRNAs and ds-oligonucleotides p3

**TABLE S3:** List of reporter or expression constructs used in this work p4

### SUPPLEMENTARY FIGURES

**Figure S1:** *Sp5* (t29291aep) single-cell expression as deduced from *Hydra* single-cell transcriptomes produced by Siebert et al (2019) and made available on Single Cell Portal; p5

**Figure S2:** Map and sequences of the *HyActin-1388*:mCherry_*HySp5-3169*:GFP construct (10'533 bp) further named *HySp5-3169*:GFP construct; p6

**Figure S3:** GFP and mCherry fluorescence in intact transgenic animals expressing the *HyActin-1388*:mCherry _*HySp5-3169*:GFP construct either in the epidermis or in the gastrodermis; p8

**Figure S4:** *GFP* expression in regenerating epidermal and gastrodermal *HySp5-3169*:GFP animals; p9

**Figure S5:** GFP and mCherry fluorescence in apical-regenerating (AR) or basal-regenerating (BR) halves of epidermal and gastrodermal *HySp5*-3169:GFP transgenic animals; p10

**Figure S6:** Impact of alsterpaullone treatment on *Sp5* and *Wnt3* expression in *Hv_AEP2* animals; p11

**Figure S7:** Impact of ALP treatment on Sp5, GFP and Wnt3 expression in epidermal and gastrodermal HySp5-3169:GFP transgenic animals; p12

**Figure S8:** Impact of ALP treatment on *GFP, Wnt3* and *Sp5* expression pattern in epidermal and gastrodermal *HyWnt3-2149*:GFP transgenic animals; p13

**Figure S9:** Impact of *β-catenin*(RNAi) on *β-catenin*, *Sp5* and *GFP* transcript levels in epidermal and gastrodermal *HySp5-3169*:GFP transgenic animals; p14

**Figure S10:** Impact of *β-catenin*(RNAi) on pseudo-bud formation and *Sp5* expression in *Hv-Basel* animals p15

**Figure S11:** Impact of *β-catenin*(RNAi) on GFP and mCherry fluorescence in epidermal and gastrodermal *HySp5-3169*:GFP transgenic animals; p16

**Figure S12:** Impact of *β-catenin*(RNAi) on GFP and mCherry immuno-detected patterns in epidermal and gastrodermal *HySp5-3169*:GFP transgenic animals; p17

**Figure S13:** Impact of *Sp5*(RNAi) on *Sp5, GFP* and *Wnt3* expression levels in epidermal and gastrodermal *HySp5-3169*:GFP transgenic animals; p18

**Figure S14:** Impact of *Sp5*(RNAi) on *GFP* expression in epidermal and gastrodermal *HySp5-3169*:GFP transgenic animals; p19

**Figure S15:** Impact of *Sp5 (RNAi)* on GFP and mCherry fluorescence in epidermal or gastrodermal *HySp5-3169*:GFP transgenic animals; p20

**Figure S16:** Impact of *Sp5(RNAi)* on GFP fluorescence in epidermal and gastrodermal *HySp5-3169*:GFP animals p21

**Figure S17:** ChIP-qPCR analysis of the Sp5-binding sites in the *HySp5* promoter using anti *HySp5* antibodies; p22

**Figure S18:** Mapping the putative *Sp5* Transcriptional Start Sites (TSS); p23

**Figure S19:** Putative *Zic4*-binding sites in the *Wnt3* and *Zic4* genomic sequences from *Hm-105 Hydra* p24

### SUPPLEMENTARY TABLES

#### TABLE S1: Accession numbers (AN) of the *Hydra* genes used in this study

| Gene name | Hydratlas.unige (*Hv_Jussy, Hv_AEP1, Hol_CS,_CR*) | UNIPROT / NCBI (*H. vulgaris, Hm-105*) | single cell portal (*Hv_AEP2*) | Genomic DNA (*Hm-105*) |
|---|---|---|---|---|
| ***β-actin*** | seq16839_loc07321 (*Hv_Jussy*)<br>c25578_g1_i02 (*Hv_AEP1*)<br>S040281c2g2_i01 (*Hol_CS*)<br>R040833c2g5_i01 (*Hol_CR*) | P17126_HYDVU<br>XP_002154696.1 | t11116aep | Sc4wPfr_135 |
| ***β-catenin*** | seq73524_loc24304 (*Hv_Jussy*)<br>c12650_g1_i01 (*Hv_AEP1*)<br>S042301c1g1_i01 (*Hol_CS*)<br>R029792c0g1_i01 (*Hol_CR*) | T2M8K9_HYDVU<br>XP_012562750.1 | t13357aep | Sc4wPfr_177 |
| ***Sp5*** | seq62049_loc21222 (*Hv_Jussy*)<br>c16537_g1 (*Hv_AEP1*)<br>S034511c0g1_i01 (*Hol_CS*)<br>R029443c0g1_i01 (*Hol_CR*) | T2MC68_HYDVU<br>XM_004206770.3 | t29291aep | Sc4wPfr_224.1 |
| ***TBP*** | seq65236_loc22074 (*Hv_Jussy*)<br>c16197_g1_i01 (*Hv_AEP1*)<br>S023957c0g1_i01 (*Hol_CS*)<br>R001357c0g1_i01 (*Hol_CR*) | T2MG73_HYDVU<br>XP_002160884.2 | t38685aep | Sc4wPfr_29 |
| ***TCF*** | seq62874_loc21437(*Hv_Jussy*)<br>c21328_g1_i01(*Hv_AEP1*)<br>S036398c0g1_i05 (*Hol_CS*)<br>R035844c0g2_i01 (*Hol_CR*) | Q9GTK1_HYDVU<br>NP_001296662.1 | t11826aep | Sc4wPfr_319 |
| ***Wnt3*** | *seq49770*_loc17775 (*Hv_Jussy*)<br>c3776_g1_i01 (*Hv_AEP1*)<br>S029978c0g1_i01 (*Hol_CS*)<br>R033453c0g1_i01 (*Hol_CR*) | Q9GTJ9_HYDVU<br>NP_001274292.1 | t14194aep | Sc4wPfr_399 |
| ***Wnt5a*** | *seq29395_loc11578* (*Hv_Jussy*)<br>*c19333_g1_i01* (*Hv_AEP1*)<br>*S037476c0g1_i02* (*Hol_CS*)<br>*R009822c0g1_i01* (*Hol_CR*) | A0A8B6XQ36_HYDVU<br>NP_001296688.1 | t21554aep | Sc4wPfr_287.2 |
| ***Wnt8*** | *seq25361_loc10235* (Hv_Jussy)<br>*c12431_g1_i01* (Hv_AEP1)<br>*S039485c1g1_i01* (Hol_CS)<br>*R036877c0g1_i01* (Hol_CR) | A0A8B7DSN5_HYDVU<br>XP_047127276.1 | t23521aep | Sc4wPfr_440 |
| ***Zic4*** | seq19466_loc08275 (*Hv_Jussy*)<br>c11132_g1_i01 (*Hv_AEP1*)<br>S030492c0g1_i01 (*Hol_CS*)<br>R028263c0g1_i01 (*Hol_CR*) | T2M6U0_HYDVU<br>XP_002153782.2 | t20709aep | Sc4wPfr_237.2 |

### TABLE S2: List of primers, siRNAs and ds-oligonucleotides

| | | |
|---|---|---|
| **Cloning primers** | HySp5 promoter Forward | CCGGATATCCTAGTTCTAATTTAGCTCTATTACGTTCGC |
| | HySp5 promoter Reverse | AACCCCTTATCAAAGAAGCCACCGGTCTAG |
| **siRNAs** | HySp5 siRNA-1 | UUA ACG AGC ACC ACA UAA A |
| | HySp5 siRNA-2 | CUA CAA CAU CCC ACA UAU A |
| | HySp5 siRNA-3 | GCA GCA CGU AUG UCA UAU U |
| | β-catenin siRNA-1 | UCA ACC UAA CAG ACA ACA A |
| | β-catenin siRNA-2 | UGA GGA GCU AUA CUU AUG A |
| | β-catenin siRNA-3 | ACG ACU CUC UGU UGA AUU U |
| | Scramble siRNA | AGG UAG UGU AAU CGC CUU G |
| **qPCR primers** | HySp5 Forward | CCAGGGTGCGGAAAGGTT |
| | HySp5 Reverse | CCAGCATGCCATCTTAAATGAG |
| | HyWnt3 Forward | GAGTTGACGGTTGCGAACTT |
| | HyWnt3 Reverse | ACATGAAACCTTGCAACACCA |
| | β-catenin Forward | TACGCAATGTTGTTGGTGCT |
| | β-catenin Reverse | GCTTCAATTCGATGGCCTAA |
| | GFP Forward | TGGAAGCGTTCAACTAGCAG |
| | GFP Reverse | AAAGGGCAGATTGTGTGGAC |
| | TBP Forward | AAGCGATTTGCAGCAGTTAT |
| | TBP Reverse | GCTCTTCACTTTTTGCTCCA |
| **qPCR primers for ChIP** | Sp5prom_F_1 | TAAGCTGTCTCCATTTCAACCA |
| | Sp5prom_R_1 | AATATTTGTTAAGTGTTTTTCGTTGG |
| | Sp5prom_F_2 | AATTGCGGTAAAGATCAGTAAGAA |
| | Sp5prom_R_2 | TGGTTGAAATGGAGACAGCTT |
| | Sp5prom_F_3 | AAGTATCAAGTTTAAAAATTCCTTCG |
| | Sp5prom_R_3 | ATTTAGAATTCTTACTGATCTTTACCG |
| | Sp5prom_F_4 | TATCTTTTCCGCCTTACGTATTC |
| | Sp5prom_R_4 | ACTGAGAAATGGCGCGTTG |
| | Sp5prom_F_5 | CAGAGAAAATATGATCGCAACG |
| | Sp5prom_R_5 | GAAACCGCCATCTTATCTTAAA |
| | Sp5prom_F_6 | AACCAAATATTTAAAATGATAAACTGG |
| | Sp5prom_R_6 | CAAAGGCGGAGTAATTAGGTG |
| | Sp5prom_F_7 | TGATTTGAAGTCAAAAACAAAATAACA |
| | Sp5prom_R_7 | TGGTTAAAAGATATAAAACGCTATTTG |
| | Sp5prom_F_8 | TGGTAAAGTTCGTAAAACCAATGA |
| | Sp5prom_R_8 | AAACATTCCGACAATCCACAG |
| | Sp5prom_F_9 | AAGTAGCGACAGCGCCAGT |
| | Sp5prom_R_9 | ATATCCTAGCCAAAACAAAACAA |
| | Sp5prom_F_10 | GGTCAGCGAGTTTGGATCAT |
| | Sp5prom_R_10 | AGCCTCAGGACTTCCCATTT |
| | Sp5prom_F_11 | CGAGCGTTGCTTTGACTTTA |
| | Sp5prom_R_11 | CAATTACGGATCACCGAAGG |
| | Sp5prom_F_12 | CTCAGTGCATCCGTTCGTT |
| | Sp5prom_R_12 | TCTTGCTTGCTTACGGATGA |
| | Sp5prom_F_13 | TGAAATATTAAAAAGACGGAAGGAA |
| | Sp5prom_R_13 | TGCAGTGAAAAGCAACAAACA |
| | Sp5prom_F_14 | CGTTCGCAAAGTTGACAAAGT |
| | Sp5prom_R_14 | CAATTTTTATAAGCGTGATAAAGCAA |
| | Sp5prom_F_15 | TCAATTTCAACAAAATAAGTGTCAA |
| | Sp5prom_R_15 | TGAAGTTTCAATCCCTTTTAAACAA |
| **double-stranded oligos for EMSA** | ds-Sp5-oligo PPA wt | TATCTTTT**CCGCCT**TACGTATTCTGTTTATCA**CCGCCT**CTTAGACCATCCCATTTGTACGTAAACAGAG |
| | ds-Sp5-oligo PPA mut | TATCTTTT**CTTCCT**TACGTATTCTGTTTATCA**CTTCCT**CTTAGACCATCCCATTTGTACGTAAACAGAG |
| | ds-Sp5-oligo PPB wt | CAGAGAAAATATGATCGCAAC**GCGCCA**TTTCTCAGTCAG**AGGCGT**GACATTAACCCCTTATCAAAGAAGCCGA |
| | ds-Sp5-oligo PPB mut | CAGAGAAAATATGATCGCAAC**GTTCCA**TTTCTCAGTCAG**ATTCGT**GACATTAACCCCTTATCAAAGAAGCCGA |

### TABLE S3: List of reporter or expression constructs used in this work

| PLASMID NAME | PLASMID DESCRIPTION | REFERENCE |
|---|---|---|
| **CONSTRUCTS TO BE EXPRESSED *EX-VIVO*** | | |
| ***HySp5-2992*:Luciferase** in pGL3 | Construct where 2'992 bp *Hydra Sp5* upstream sequences directing luciferase expression in human HEK293T cells | *[1]* |
| ***HySp5-2828*:Luciferase** in pGL3 | Construct derived from the *HySp5-2992*:Luciferase plasmid after deleting 164 bp of the proximal *Hy Sp5* promoter sequences | *This work (Fig. 6, Fig. 7)* |
| ***HySp5-2992-MBS1*:Luc**<br>***HySp5-2992-MBS2*:Luc**<br>***HySp5-2992-MBS3*:Luc**<br>***HySp5-2992-MBS4*:Luc**<br>***HySp5-2992-MBS5*:Luc** | Constructs derived from the *HySp5-2992*:Luciferase construct after mutating in each one out of the 5 Sp5-binding sites located in the proximal promoter: BS1 at position -129, BS2 at position -105, BS3 at position -52, BS4 at position -34, BS5 at position +18 | *This work (Fig. 6, Fig. 7)* |
| ***CMV*:HySp5-420** in pCS2+<br>***CMV*:HySp5-337-ΔDBD** | Expression constructs designed to express either full length Sp5 (HySp5-420) or Sp5-337 lacking its DNA-binding domain (ΔDBD) | *[1]* |
| **pCS2+-HySp5-420** | bacterial vector to express the *Hydra* Sp5 protein, here full length | *[1]* |
| ***HyWnt3-2149*:Luciferase** in pGL3 | Construct where 2'149 bp *Hydra Wnt3* upstream sequences directing luciferase expression in human HEK293T cells | *[1]* |
| ***CMV*:hu_β-CateninΔ45** in pFLAG | Construct designed to express a truncated version of β-catenin protein, constitutively active in human cells | *[1,2]* |
| **TOPFLASH**<br><br>**FOPFLASH** | TOPFLASH: Reporter construct where 6 arrowed consensus TCF binding sites upstream of the minimal TK promoter direct Luciferase expression;<br>FOPFLASH: Reporter construct where 6 arrowed mutated TCF binding sites upstream of the minimal TK promoter direct Luciferase expression | *[3]* |
| **pCAG-FLAG-TCF-1** | Construct designed to express a FLAG-tagged version of the human TCF1 transcription factor in human cells | *[4]* |
| ***HyZic4-3505*:Luciferase** in pGL3 | Reporter construct where 3'505 bp of the *HyZic4* upstream sequences drive luciferase expression in human HEK293T cells | *[5]*<br>*This work (Fig. 7, Fig. S21)* |
| ***CMV*:HyZic4-431** in pCS2+<br>***CMV*:HyZic4-ΔDBD** | Expression constructs designed to express Zic4 either full length (HyZic4-431) or lacking its DNA-binding domain (HyZic4-ΔDBD) | *[5]* |
| **CONSTRUCTS TO BE EXPRESSED IN *HYDRA*** | | |
| ***HyAct-1388*:eGFP** (hoTG) | Transformation reporter construct where 1'289 bp upstream and 99 bp 5'UTR sequences of *Hydra Actin* direct ubiquitous GFP expression (full sequence available @ NCBI: DQ369740) | *[6]* |
| **hoTG-*HyWnt3FL*:eGFP-*HyAct*:dsRED** | Tandem reporter construct where 2'149 bp upstream sequences of *HyWnt3* direct eGFP expression and 1'289 bp upstream sequences of *HyActin* direct ubiquitous RFP expression | *[7]* |
| ***HyAct-1388*:mCherry:: *HySp5-3169*:eGFP** in PBSSA | Tandem reporter construct where 1'289 bp upstream and 99 bp 5'UTR sequences of *Hydra Actin* direct mCherry expression and 3'169 bp upstream sequences of *Hydra Sp5* direct eGFP expression | *This work (see full sequence in Fig. S2)* |
| **CONSTRUCTS TO DETECT GENE EXPRESSION IN *HYDRA*** | | |
| ***GFP-663**_pGEM-T-Easy* | Plasmid to produce a 663 nt-long *GFP* riboprobe | *This work* |
| ***HySp5-502**_pGEM-T-Easy* | Plasmid to produce a 502 nt-long *Sp5* riboprobe – | *(Vogg et al. 2019)* |
| ***HyWnt3-1092**_pGEM-T-Easy* | Plasmid to produce a 1092 nt-long *Wnt3* riboprobe – | *(Vogg et al. 2019)* |

### SUPPLEMENTARY FIGURES

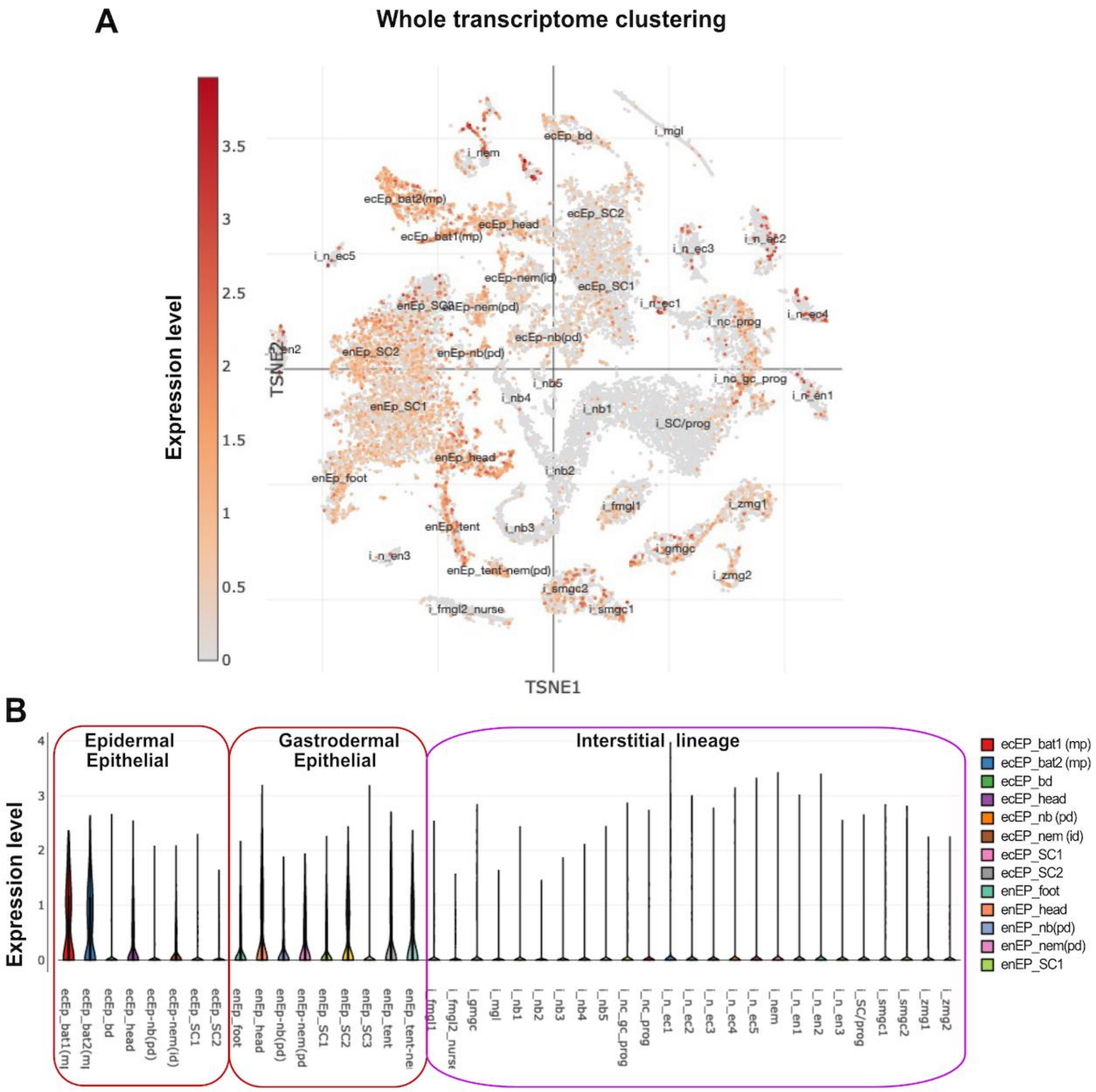


**Figure S1. *Sp5* (t29291aep) single-cell expression as deduced from *Hydra* single-cell transcriptomes produced by Siebert et al (2019) and made available on Single Cell Portal**

**(A)** tSNE plot of SNN clustered single cell data mapped to the LRv2 transcriptome reference. **(B)** Distribution of *Sp5* transcripts per cell type. The *Hydra Sp5* sequence from the *Hv_AEP* strain (c16537_g1 available on HydrAtlas) was used for a blast search on the Juliano aepLRv2 reference transcriptome. The protein sequences deduced from c16537_g1/Sp5 and t29291aep/Sp4 cDNAs are 100% identical. Note the predominant expression in the epithelial cells (Ep) from the epidermis (also called ectoderm, ecEP) or the gastrodermis (also called endoderm, enEp). Epithelial sub-populations were identified as stem cells (epEP_SC1, epEP_SC2, enEp_SC1, enEp_SC2, enEp_SC3), suspected phagocytosis doublet (ecEp-nb(pd), enEp-nb(pd), enEp-nem(pd)), epidermal cells that contain integration doublet (ecEp-nem(id)), tentacle-specific in the gastrodermis (enEp_tent, enEp_tent-nem), tentacle-specific epidermal battery cells that are multiplet (ecEP_bat1(mp), ecEP_bat2(mp)), apical-specific (ecEp_head, enEp_head), basal-specific (ecEp_bd, enEp_foot). **Abbreviations: bat:** battery cell, **bat(mp):** multiplet battery cells, **bd:** basal disc, **ec:** ectoderm (=epidermis), **en:** endoderm (=gastrodermis), **ep:** epithelial, **fmgl:** female germline, **gc:** gland cell, **gmgc:** granular mucous gland cell, **i:** cell from the interstitial lineage; **id:** integration doublet, **mgl:** male germline, **mp:** multiplet, **nb:** nematoblast, **n**: neuronal cell, **nem:** nematocyte, **pd:** suspected phagocytosis doublet, **prog:** progenitors, **SC:** Stem Cell, **smgc:** spumous mucous gland cell, **tent:** tentacle, **zmg:** zymogen gland cell.

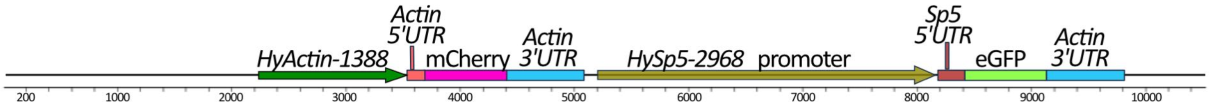


#### *pBSSA-AR plasmidic sequence (2'244 bp)*

```
   1 gtggcacttttcggggaaatgtgcgcggaacccctatttgtttatttttctaaatacattcaaatatgtatccgctcatgagacaataaccctgataaat
 101 gcttcaataatattgaaaaaggaagagtatgagtattcaacatttccgtgtcgcccttattcccttttttgcggcattttgccttcctgtttttgctcac
 201 ccagaaacgctggtgaaagtaaaagatgctgaagatcagttgggtgcacgagtgggttacatcgaactggatctcaacagcggtaagatccttgagagtt
 301 ttcgccccgaagaacgttttccaatgatgagcacttttaaagttctgctatgtggcgcggtattatcccgtattgacgccgggcaagagcaactcggtcg
 401 ccgcatacactattctcagaatgacttggttgagtactcaccagtcacagaaaagcatcttacggatggcatgacagtaagagaattatgcagtgctgcc
 501 ataaccatgagtgataacactgcggccaacttacttctgacaacgatcggaggaccgaaggagctaaccgcttttttgcacaacatgggggatcatgtaa
 601 ctcgccttgatcgttgggaaccggagctgaatgaagccataccaaacgacgagcgtgacaccacgatgcctgtagcaatggcaacaacgttgcgcaaact
 701 attaactggcgaactacttactctagcttcccggcaacaattaatagactggatggaggcggataaagttgcaggaccacttctgcgctcggcccttccg
 801 gctggctggtttattgctgataaatctggagccggtgagcgtgggtctcgcggtatcattgcagcactggggccagatggtaagccctcccgtatcgtag
 901 ttatctacacgacggggagtcaggcaactatggatgaacgaaatagacagatcgctgagataggtgcctcactgattaagcattggtaactgtcagacca
1001 agtttactcatatatactttagattgatttaaaacttcatttttaatttaaaaggatctaggtgaagatcctttttgataatctcatgaccaaaatccct
1101 taacgtgagttttcgttccactgagcgtcagaccccgtagaaaagatcaaaggatcttcttgagatcctttttttctgcgcgtaatctgctgcttgcaaa
1201 caaaaaaaccaccgctaccagcggtggtttgtttgccggatcaagagctaccaactctttttccgaaggtaactggcttcagcagagcgcagataccaaa
1301 tactgtccttctagtgtagccgtagttaggccaccacttcaagaactctgtagcaccgcctacatacctcgctctgctaatcctgttaccagtggctgct
1401 gccagtggcgataagtcgtgtcttaccgggttggactcaagacgatagttaccggataaggcgcagcggtcgggctgaacggggggttcgtgcacacagc
1501 ccagcttggagcgaacgacctacaccgaactgagatacctacagcgtgagctatgagaaagcgccacgcttcccgaagggagaaaggcggacaggtatcc
1601 ggtaagcggcagggtcggaacaggagagcgcacgagggagcttccagggggaaacgcctggtatctttatagtcctgtcgGgtttcgccacctctgactt
1701 gagcgtcgatttttgtgatgctcgtcaggggggcggagcctatggaaaaacgccagcaacgcggcctttttacggttcctggccttttgctggccttttg
1801 ctcacatgttctttcctgcgttatcccctgattctgtggataaccgtattaccgcctttgagtgagctgataccgctcgccgcagccgaacgaccgagcg
1901 cagcgagtcagtgagcgaggaagcggaagagcgcccaatacgcaaaccgcctctccccgcgcgttggccgattcattaatgcagctggcacgacaggttt
2001 cccgactggaaagcgggcagtgagcgcaacgcaattaatgtgagttagctcactcattaggcaccccaggctttacactttatgcttccggctcgtatgt
2101 tgtgtggaattgtgagcggataacaatttcacacaggaaacagctatgaccatgattacgccaagcgcgcaattaaccctcactaaagggaacaaaagct
2201 ggtaccgggccccccctcgaggtcgactctagaggatccccccat             2244
```

#### *Actin promoter sequences (1'289 bp) + Actin 5'UTR (99 bp) + Actin coding sequence (27 bp) + MCS (27 bp)*

```
-1289 cgatctgactaacctaaccagtgcaaaaaaatttaaaagatttgcattgtgaaagttagaatattataaaaaatctaaaacgagtattactcgagtaaat
-1189 gttatacgatctatagattaaatatattaaaaatgtatagcgaatgttaaactaaatatataatataaacttgaaaacttactaaattgcaaaaactcaa
-1089 aaccgactgtatcattttacaggaaaccgttattcaagatacttaagttgtttactacattattataacatcttgcaattagcaagacaatcgttattt
 -989 taacatcacggtatcgaaaggattttgagaaattttattgaaacattttaaacaaaaaatatcatatttagatgcattttaagccgagatgcaggattct
 -889 gaatgaaaaagaaaaaaagaagtctcggtagagtaaaagtgatcggtttgcaactgtaaaatttattgaagtaccaataattttatttaaaataaaactg
 -789 aaatataaagttaaagttgctgttctataagtttaccgaattttaaaaccattgtaacgctagagtaatatttgagtctactaagttagtccccgcactt
 -689 tttaatcaagcaataaatacccaaactttgcttattcaaatcaataaaccaatatatctcttaaaataaagtaaaaacttctgaaattctataaaaaaaa
 -589 atttaatttcgaaatatcaaatgtaacttcaacaccgcactattttcttttaaacaactgatatagtaattacttctcaaaaacgttatctcaaggtttg
 -489 tgatgtacttaaaaccactcctattttgttacgcgtttaaaaaagcaaacataagttggtttctattgatgaatgagaacatatttcatttaaagttaaa
 -389 atcctaccagtggtttcactgtacgtaaacaccgtcaaaaaaacaggaacgtttttaaagattaataattgaagtaaaaaaaatttaataccggggtta
 -289 aaaaaatcttttaaaataattataaatatatatattaaaatttataaattttaaacacatttaaaatatatattaagtataataaaagtaatattataa
 -189 aaaaaaatttaatttataattattttattaaatttataaataataggtaaaacttacatatccgtttattttttcttaataaaataacgcgtgcaaa
                                                                                                   TSS +1
  -89 ttttgtccatataaagaccttttcgaacaataactttttgcttagccgtttttttttcttatatggtcaaaaaagcgctcaagcgattCAccataaaaa
                                                                                                 99
   12 gcgcaattagttcagcgttcgttattcagaagcttcagctttgcttgatactcagctcttctctttttaaacaaaacacttaatcaaaATGGCCGATGAT
                                                                                            M  A  D  D
      GAAGTTGCCGCCCTC GCTGCAGCCCCGGTAGAAAAAGTCGAC
      E  V  A  A  L
```

#### *mCherry coding sequence (711 bp) + MCS (14 bp)*

```
  1 ATGGTTAGTAAAGGAGAAGAAGATAATATGGCTATTATTAAAGAATTTATGAGATTTAAAGTTCATATGGAAGGATCTGTTAATGGTCATGAATTTGAAA
101 TTGAAGGTGAAGGAGAAGGTAGACCATATGAAGGTACTCAAACAGCTAAACTTAAAGTTACTAAAGGTGGACCATTACCTTTTGCATGGGATATTCTTTC
201 ACCACAATTTATGTATGGAAGTAAAGCTTATGTTAAACATCCAGCAGATATTCCTGATTATCTTAAACTTTCATTTCCTGAAGGTTTTAAATGGGAAAGA
301 GTTATGAATTTTGAAGATGGTGGAGTTGTTACTGTTACACAAGATTCTTCATTACAAGATGGAGAATTTATTTATAAAGTTAAACTTAGAGGAACTAATT
401 TTCCAAGTGATGGTCCTGTTATGCAAAAGAAAACTATGGGTTGGGAAGCTAGTTCTGAAAGAATGTATCCTGAAGATGGAGCACTTAAAGGTGAAATTAA
501 ACAAAGACTTAAACTTAAAGATGGTGGACATTATGATGCTGAAGTTAAAACTACATATAAAGCTAAAAAACCAGTTCAATTACCTGGAGCTTATAATGTT
601 AATATTAAACTTGATATTACTTCTCATAATGAAGATTATACAATTGTTGAACAATATGAAAGAGCAGAAGGTAGACATTCAACAGGTGGAATGGATGAAT
701 TATATAAATAA        CATTCGTAGAATTC
```

#### *Actin 3' UTR (677 bp) + pBSSA-AR plasmidic sequence (110 bp)*

```
  1 acaattcgattatatttatactggactatttttacatctgttcggttattttcacatttatttttctatatatatcttataaacgttttaaaacccatgt
101 aattttgttaagctgtaatataaaagacgtcctaacaaacttcttttattactgaatttcctttaattataataaataacaagttttaaaataaattca
201 ggcaattaaggcgctcctgaggtactaaaattaatgtaaacatttaaaattaacttggatggtcttaagtactgtactcgtgatttttgttatactttatt
301 attagaaaagtcgtctattaacttttgttccttaatttacttgattaaattgtcgcttaatttatcaaatcaggttttgcgcgttattttagagaaaaa
401 cttattagaaaaatgaataagcaaagtttaggctaacatgttttttttattattttaaatagttcaagtcaatgacgtataaaatgcatttgcaaaaaatt
501 ttaagtaaccctataaacttagcaatagtagatactggatgcaagcattcagtagcagcattgcatatctgctgtctttacgtacaaataacagcaaaaa
601 tggacctttattggcttcacatcgtcgtaaaacatgtgttattggacttgtcacaaatgtgttaagtatacagagct            677
  1 tagctcttgatgttgatcactagttctagagcggccgccaccgcggtggagctcggcgcgccggtaccgggccccccctcgaggtcgacggtatcgataa
101 gcttgatatc
```

#### *Hydra Sp5 promoter (2'968 bp) + Sp5 5'UTR (201 bp) + Sp5 coding sequence (27 bp) + MCS (20 bp)*

```
-2968  ctagttctaatttagctctattacgttcgcaaagttgacaaagtcgcaaatttttttctttttcaaaagacctcccattcattttaataaagactggttc
-2868  taattttgctttatcacgcttataaaaattgacaaagtcgcaagtactttgtttttttaaaagacctcccattcattgagataaatactagttattttcat
-2768  gtatcattgaaatagtaaacaattcattctagtttttatttgtctagggcagtatttcacaccttccacaagtgcgaaacgttttatttacttgattgag
-2668  taaattaatttaaaataaaataaaaaaataagaacttaatgtgaaaaaaaaaacaaaacaaaaacaataaaaaaaaaaacaatgtaaaatatttctcata
-2568  gctgttgaaatattaaaaagacggaaggaaaaatataacggcaaagctaaattcttttctgcgtattgcTTTTTATctctttattgtctgtttgttgctt
-2468  ttcactgcattctattttgtgcttaataaatctcaatcgattttaaggaatgactaggatgtttcattttgtatatatcaataactgaaatattaaaaat
```

```
-2368  CTCCTCAGTGCATCCGttcgttagacaattggggtattaactcaattatttcctgaatataaactcaacaagtaaaaaagtttcatccgtaagcaagca
-2268  agaataacgacacttgtttacatttaagaatttcttaactttatgtaaaaaacaattcttagttaaaaacgaagtaaaagggttttaatttttttgttttt
-2168  tagttgaaaacaaattgctaaataaaacttaatttaaaaaaaaaaaaaccaaatatttaaaatgataaactggttaaaattaatagattatacaaaccatt
                                                                                    Sp5-BS
-2068  gtaagcatttaaaaacaattttttttttataaaaacaacaacaaaaaaatttcacctaattactCCGCCTttgttaataaaacctccgttttacagttaa
-1968  aagtaatgtatgaaccgtaatccctattacaaaagaaatggtatattgtttataaagacgttttgatgattgtagcttattttatattttgttgctttt
-1868  gtttgttttccttgttgttgtttttaatttgattagttatattcttacggccttacgaccacgtttgctgttttaaatcgcgagcgttgctttgacttta
-1768  cacgaagcctctaaaacaaacataaagagaattcatcgaaagtaaaaaaaatcatgctgaccttcggtgatccgtaattgaaattgatatatattttctc
-1668  cctattttgacatataaatggttaaagttaatctttttattaaactcaatcaatttcaacaaaataagtgtcaagtttactgcttatttcaagtaacaaa
-1568  taagtcttattgtaaataaagttattgtttaaaagggattgaacttcaatcattattgtaataaagaagaatttcatgtaagatttgtatttattaaaa
-1468  ttaaataaactaaatgaacaaaagcgctacgtcaaatttatattttgattattaagaggaatttttttacctaatgtaaaactactgtaaaaatcgact
-1368  gaatcaataaggtcagagagactaggtcagcgagtttggatcattaaaatcgataacaataattaacgatatagtttataatgataggaaacttacactt
                                                                                                    Sp5-BS
-1268  gacatttaaatgggaagtcctgaggctataacgttcgtttgtcgtgggtagataagccaattgacaaaaccatcatcttatatttttatggGCGCCAaat
                         TCF-BS
-1168  gtttatcatgtttaatttCTTTTATataaatgataaaaacatttaaccacaaattatttttttttatctccaaatgaaatcaagaacttttaagtcataaa
                  Sp5-BS
-1068  aagtggcgacaGCGCCAgtgataatcatagacaagtgtacacattagcttatcaaaaagtacgctagagtaaagcttattgttttgttttggctaggata
                                    TCF-BS
 -968  tatccttctcgttaaaataatttgtcctaCTTTTATatacgatattcatatttttaggtttcttgtttcgatatatatatatatttcgttatgtttgtat
 -868  gtatatatgtgtttgtttgtatgtgtatgtaaaaatatgaaatatactttttgcaaatctttgtagaagtttaataaataaagtatcaagtttaaaaatt
         TCF-BS
 -768  cCTTCGATattttaaagcttcaatttggttgggcgtagacacattagtaattgcggtaaagatcagtaagaattctaaatagacgttaattttaaAACC
        Zic-BS     TCF-BS
 -668  TGGCCTGCCCCTTGATtatttaatttgaaatttttaagctgtctccatttcaaccacagtatcaatGGGTCGGCaaaaaaagaaagattgaacgttttat
 -568  caattttaccaacgaaaaacacttaacaaatattgtagtactttttaagtttaaatgtttttgtaaaactgttatttttaaaataaacctttttacttc
                                                                                                Zic-BS
 -468  tttttttttttttgaaatcgttttaaactgatatttaataaaagcttaaatataaatgtggtaaagttcgtaaaaccaatgaaGGCAGGTGCCGGCat
                                                                                         TCF-BS
 -368  agatgaaagtgaaagaacaattttttttttattgaacttcacatttactgtggattgtcggaatgttttactattaagttgATTTGAAGTcaaaaacaaaa
                                                                                   TSS3 -194  (H. oligactis)
 -268  taacaaaatcaaccaatgaacttccttagaaattgtttaatcataaaccaatcaaatagcgtttTATAtctttt_A_accaataagataacaattttattgt
                       TSS2 -151  (Hm105)         Sp5-BS1                   Sp5-BS2
 -168  tttgtcggcatcttaag_A_tattaaaagttaatatcttttCCGCCTtacgtattctgtttatcaCCGCCTcttagaccatcccatttgtacgtaaacagag  PP4/PP5
                          Sp5-BS3          Sp5-BS4              TCF-BS       TSS1 (H. AEP)        Sp5-BS5
  -68  aaaatatgatcgcaacGCGCCAtttctcagtcagAGGCGTgacattaacccccttATCAAAGaagccgaAgctatttaagataagaTGGCGGtttcttctg  PP4/PP5
  +33  atgtctgaatgaattgtttcggtttttttttgaagtaaggataacgttaagaacgatttgaaatgcccagctgatttaaatagaattgaagtagaaagcat
                                                                                    +202
 +133  tcagttaaactttctactttaaagtgcaaagacgttcataagaaatacatttcagtaaaattaagtaaaATGTCACCTCCAAGTCGTGTTCCAA +227
                                                                                    M  S  P  P  S  R  V  P
       CTGCAGCACCCGGGGAAAAA
```

***eGFP coding sequence (717 bp)*** ***+ MCS (14 bp)***

```
  1.ATGAGTAAAGGAGAAGAACTTTTCACTGGAGTTGTCCCAATTCTTGTTGAATTAGATGGTGATGTTAATGGGCACAAATTTTCTGTCAGTGGAGAGGGTG
101 AAGGTGATGCAACATACGGAAAACTTACCCTTAAATTTATTTGCACTACTGGAAAACTACCTGTTCCATGGCCAACACTTGTCACTACTTTCTGTTATGG
201 TGTTCAATGCTTTTCAAGATACCCAGATCATATGAAACGGCATGACTTTTTCAAGAGTGCCATGCCCGAAGGTTATGTACAGGAAAGAACTATATTTTTC
301 AAAGATGACGGGAACTACAAGACACGTGCTGAAGTCAAGTTTGAAGGTGATACCCTTGTTAATAGAATCGAGTTAAAAGGTATTGATTTTAAAGAAGATG
401 GAAACATTCTTGGACACAAATTGGAATACAACTATAACTCACACAATGTATACATCATGGCAGACAAACAAAAGAATGGAATCAAAGTTAACTTCAAAAT
501 TAGACACAACATTGAAGATGGAAGCGTTCAACTAGCAGACCATTATCAACAAAATACTCCAATTGGCGATGGCCCTGTCCTTTTACCAGACAACCATTAC
601 CTGTCCACACAATCTGCCCTTTCGAAAGATCCCAACGAAAAGAGAGACCACATGGTCCTTCTTGAGTTTGTAACAGCTGCTGGGATTACACATGGCATGG
701 ATGAACTATACAAATAG    CATTCGTAGAATTC
```

***Actin 3' UTR (677 bp)***

```
  1 acaattcgattatatttatactggactatttttacatctgttcggttattttcacatttatttttctatatatatcttataaacgttttaaaacccatgt
101 aatttttgttaagctgtaatataaaagacgtcctaacaaacttcttttattactgaatttcctttaattataataaataacaagttttaaaataaattca
201 ggcaattaaggcgctcctgaggtactaaaattaatgtaaacatttaaaattaacttggatggtcttaagtactgtactcgtgatttgttatactttatt
301 attagaaaagtcgtctattaacttttgttccttaatttacttgattaaattgtcgcttaatttatcaaatcaggttttgcgcgttattttagagaaaaa
401 cttattagaaaaatgaataagcaaagtttaggctaacatgtttttttattattttaaatagttcaagtcaatgacgtataaaatgcatttgcaaaaaatt
501 ttaagtaaccctataaacttagcaatagtagatactggatgcaagcattcagtagcagcattgcatatctgctgtctttacgtacaaataacagcaaaaa
601 tggacctttattggcttcacatcgtcgtaaaacatgtgttattggacttgtcacaaatgtgttaagtatacagagct      677
```

***pBSSA-AR plasmidic sequence (713 bp)***

```
    1.......10........20........30........40........50........60........70........80........90........1
  1 tagctcttgatgttgatcactagagcggccgccaccgcggtggagctcggcgcgccagctccaattcgccctatagtgagtcgtattacgcgcgctcact
101 ggccgtcgttttacaacgtcgtgactgggaaaaccctggcgttacccaacttaatcgccttgcagcacatccccctttcgccagctggcgtaatagcgaa
201 gaggcccgcaccgatcgcccttcccaacagttgcgcagcctgaatggcgaatggaaattgtaagcgttaatattttgttaaaattcgcgttaaatttttg
301 ttaaatcagctcattttttaaccaataggccgaaatcggcaaaatcccttataaatcaaaagaatagaccgagatagggttgagtgttgttccagtttgg
401 aacaagagtccactattaaagaacgtggactccaacgtcaaagggcgaaaaaccgtctatcagggcgatggcccactacgtgaaccatcaccctaatcaa
501 gtttttggggtcgaggtgccgtaaagcactaaatcggaaccctaaagggagcccccgatttagagcttgacggggaaagccggcgaacgtggcgagaaa
601 ggaagggaagaaagcgaaaggagcgggcgctagggcgctggcaagtgtagcggtcacgctgcgcgtaaccaccacacccgccgcgcttaatgcgccgcta
701 cagggcgcgtcag      713
```

**Figure S2. Map and sequences of the *HyActin-1388*:mCherry_*HySp5-3169*:GFP construct (10'533 bp) further named *HySp5-3169*:GFP construct**

The *Sp5* upstream sequences are composed of 201 bp 5' untranslated (UTR) and 2968 bp promoter sequences.

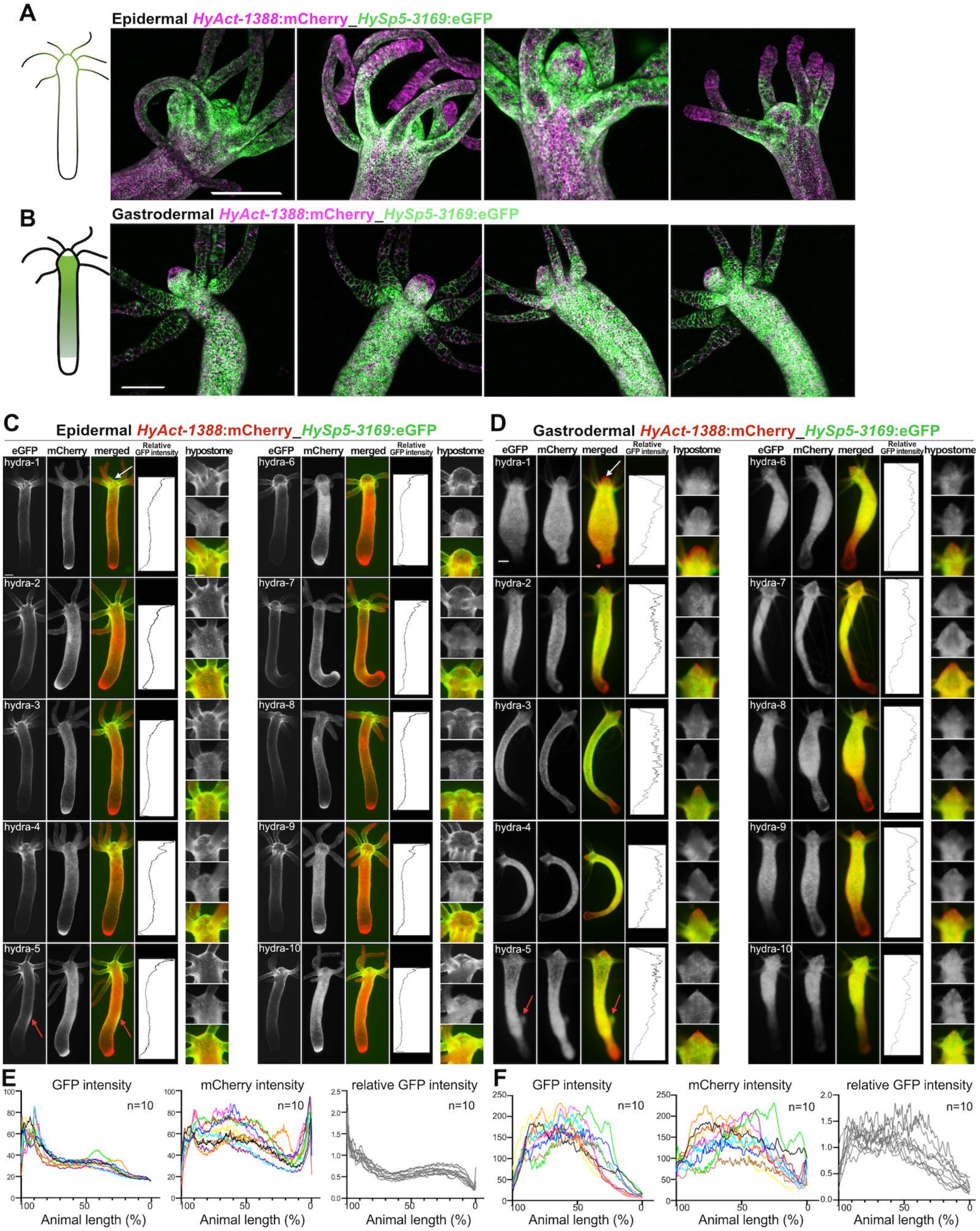


**Figure S3. GFP and mCherry fluorescence in intact transgenic animals expressing the *HyActin-1388*:mCherry _*HySp5-3169*:GFP construct either in the epidermis or in the gastrodermis**

**(A-D)** Live imaging of GFP (green) and mCherry (magenta or red) fluorescence in transgenic animals constitutively expressing the *HyActin*-1388:mCherry_*HySp5-3169*:GFP construct in their epidermal (A, C) or gastrodermal (B, D) epithelial cells. In panels C and D, magnified views of the apical region are shown on the right, white arrows indicate the most apical region, which does not express gastrodermal GFP fluorescence, red arrowhead the basal region and red arrows the GFP-positive budding regions. **(E, F)** Graphs displaying GFP (left), mCherry (middle) and relative GFP (right) fluorescence intensities along the apical (100%) to basal (0%) axis of the animals shown in panels C and D. The relative GFP intensity is calculated from the ratio between the GFP and mCherry fluorescence. Supplement to Figure 1E, 1F, 1I, 1K. Scale bars: 250 µm in all panels.

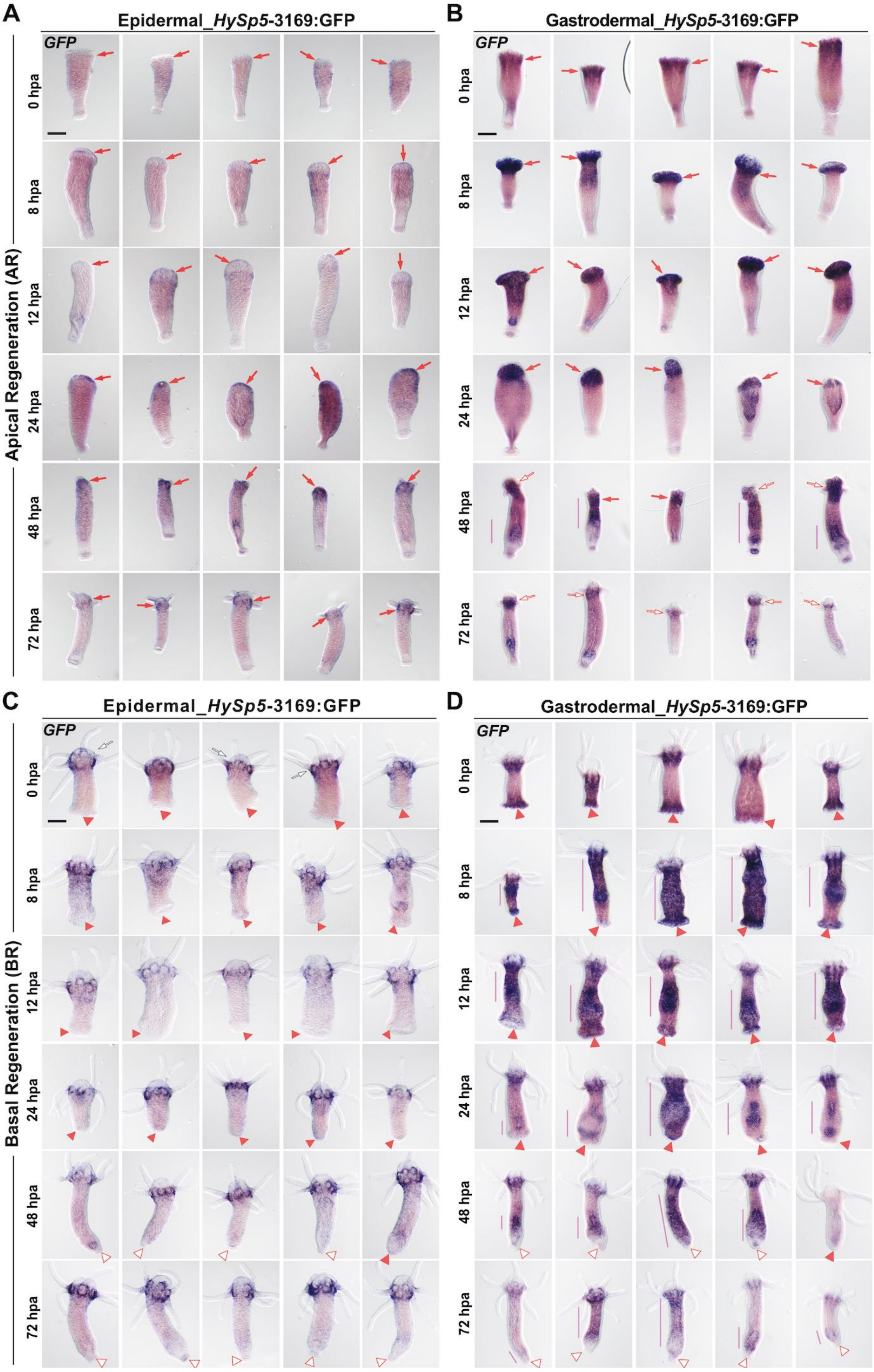


**Figure S4.** ***GFP*** **expression in regenerating epidermal and gastrodermal** ***HySp5-3169*****:GFP transgenic animals**
*GFP* expression in lower halves that are apical-regenerating (AR) **(A, B)** and upper halves that are basal-regenerating (BR) **(C, D)** from epidermal_ **(A, C)** and gastrodermal_ **(B, D)** *HySp5-3169*:GFP transgenic animals taken at 0, 8, 12, 24, 48 and 72 hours post-amputation (hpa). Vertical bars indicate areas of *GFP* expression along the body column. White arrows point to original heads; red arrows to apical-regenerating tips; white arrows outlined red to regenerated heads; white triangles to original basal discs; red triangles to basal-regenerating tips, and white triangles outlined red to regenerated basal discs. Scale bar: 250 µm. **Supplement to Figure 2A, 2B.**

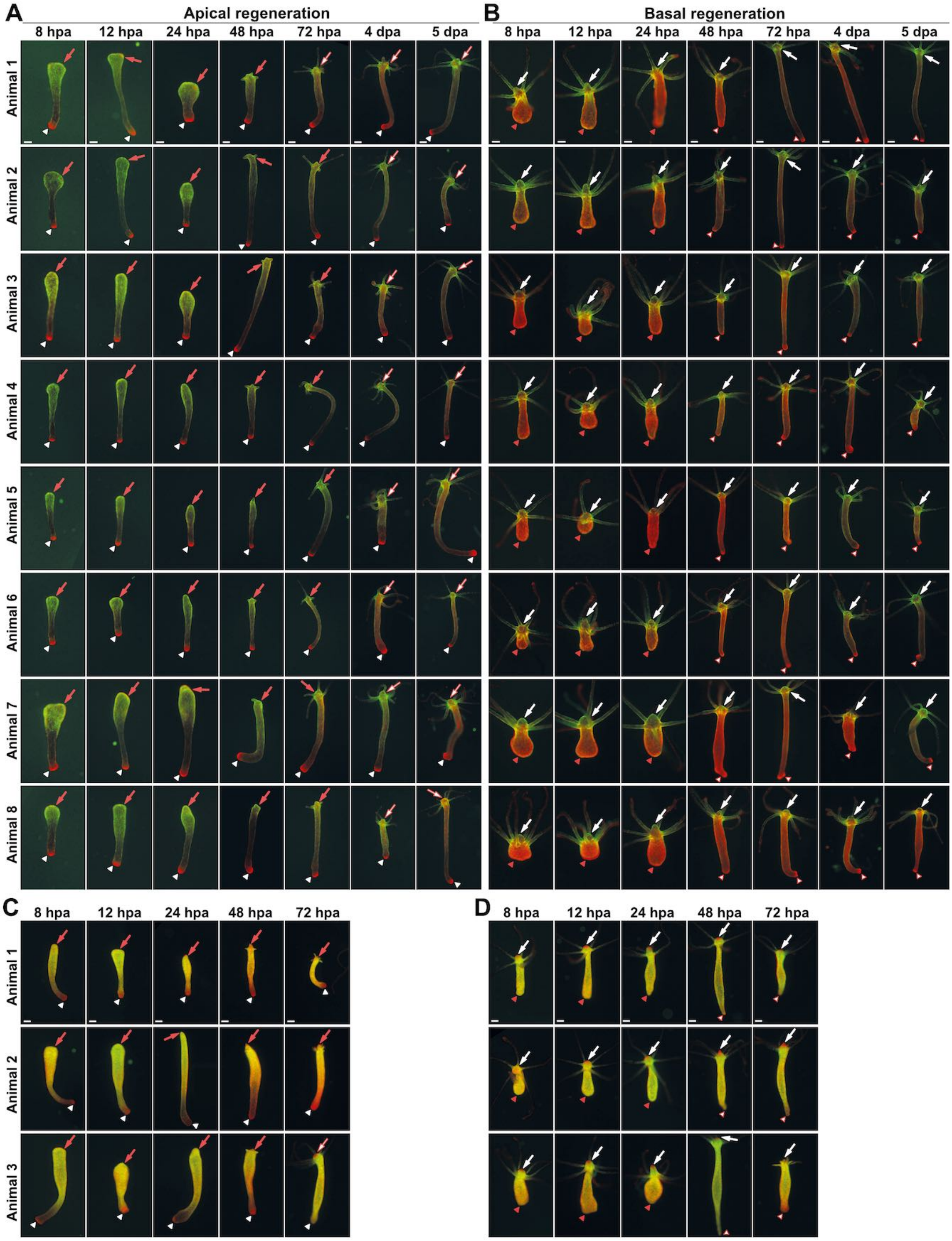


**Figure S5. GFP and mCherry fluorescence in apical-regenerating (AR) or basal-regenerating (BR) halves of epidermal and gastrodermal *HySp5*-3169:GFP transgenic animals**

Live imaging of GFP (green) and mCherry (red) fluorescence in lower (A, C) and upper (B, D) halves of epidermal_ **(A, B)** and gastrodermal_ **(C, D)** *HySp5-3169*:GFP animals pictured at indicated time-points; hpa: hours post-amputation, dpa: days post-amputation. White arrows point to original heads; red arrows to AR tips; white arrows outlined red to regenerated heads; white arrowheads to original basal discs; red arrowheads to BR tips; white arrowheads outlined red to regenerated basal discs. Scale bar: 250 µm. **Supplement to Figure 2C and 2D.**

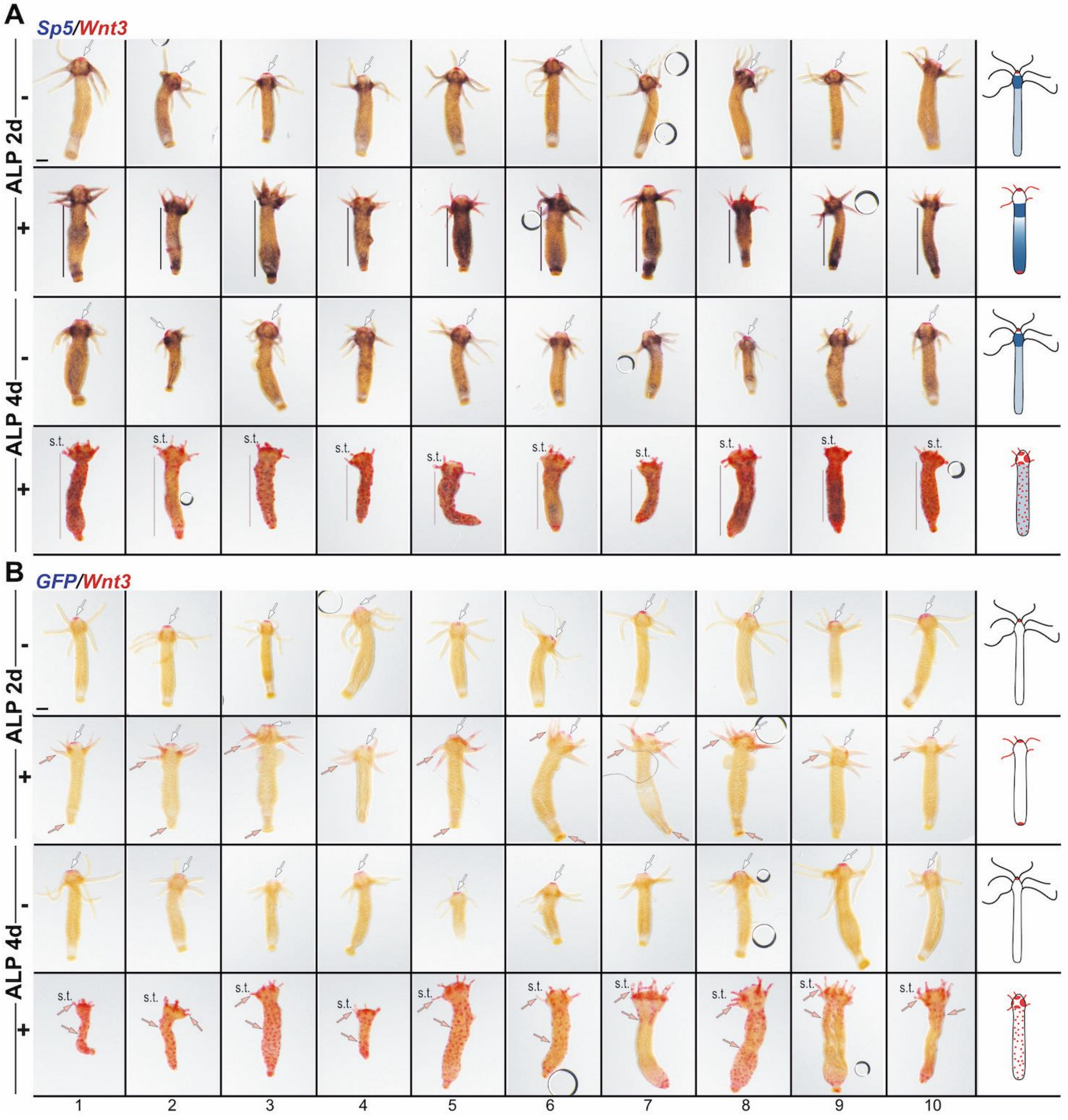


**Figure S6. Impact of alsterpaullone (ALP) treatment on *Sp5* and *Wnt3* expression in *Hv_AEP2* animals**
Co-detection of *Sp5* (dark blue) and *Wnt3* (red) **(A)**, or of *GFP* (dark blue) and *Wnt3* (red) **(B)** in wild-type *Hv_AEP2* animals left untreated (DMSO) or exposed to alsterpaullone (ALP) for 2 or 4 days. White arrows point to apical *Wnt3* homeostatic expression; salmon arrows to ectopic *Wnt3* expression; vertical bars to *Sp5* expression along the body column; s.t.: short tentacles. As expected, non-transgenic *Hv_AEP2* animals do not express *GFP*. Schematic views of *Hydra* polyps on the right depict the typical expression profile of each condition. Scale bar: 200 µm. **Supplement to Figure 3B.**

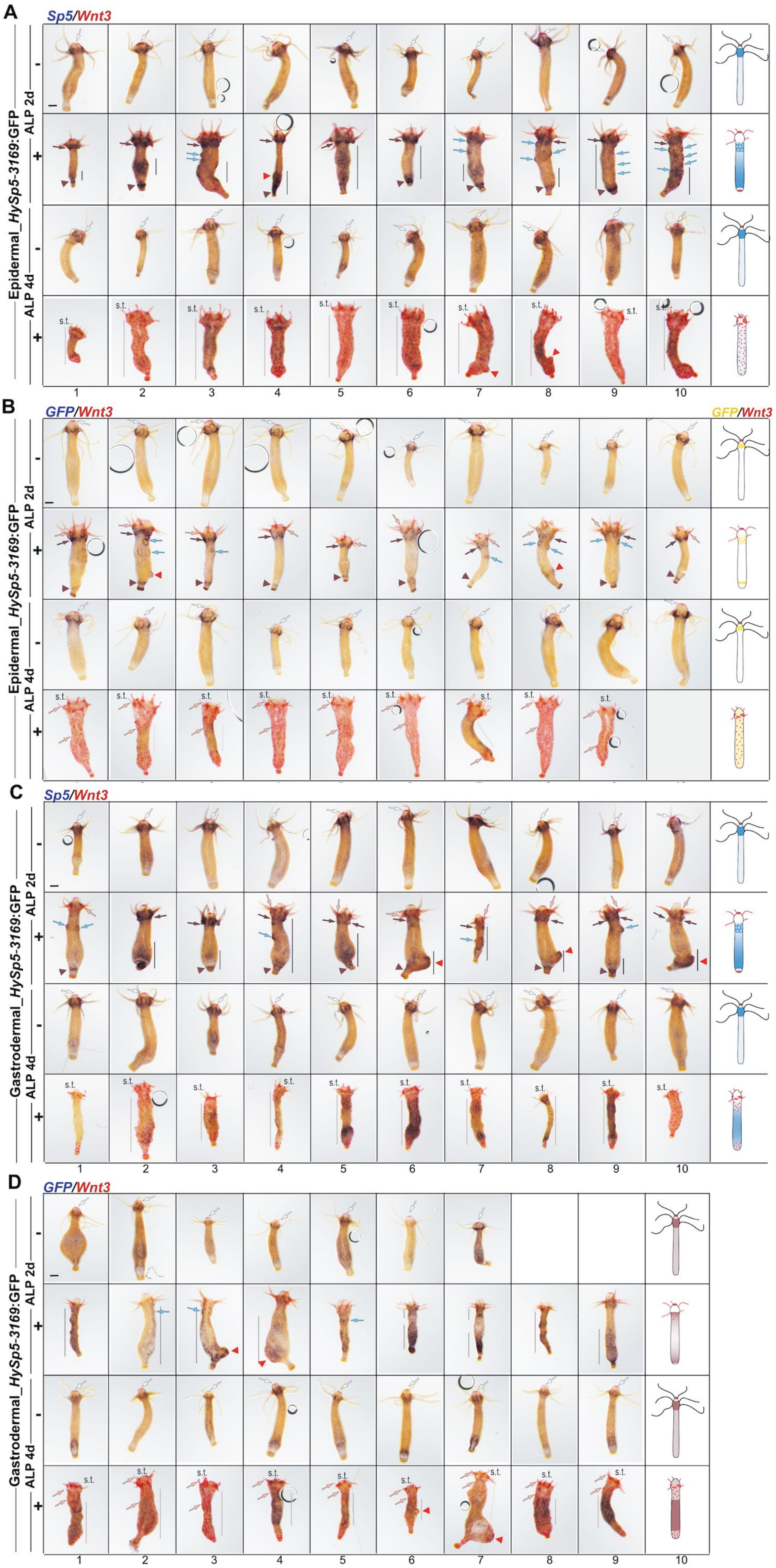


**Figure S7. Impact of ALP treatment on *Sp5, GFP* and *Wnt3* expression in epidermal and gastrodermal *HySp5-3169:GFP* transgenic animals.**

Co-detection of *Sp5* (dark blue) and *Wnt3* (red) expression **(A, C)** or of *GFP* (dark blue) and *Wnt3* (red) **(B, D)** in animals treated with DMSO (-) or ALP (+) for 2 or 4 days. In all panels brown arrows point to *Sp5* (A, C) or *GFP* (B, D) expression in the upper body column immediately below the tentacle ring; light blue arrows to spots of *Sp5* (A, C) or *GFP* (B, D) expression in the body column; vertical bars to diffuse *Sp5* or *GFP* expression along the body column; black triangles to *Sp5* (A) or *GFP* (B) basal expression; red triangles indicate pseudo-bud structures growing from the lower body column expressing *Sp5* (A, C) or *GFP* (B, D); white arrows to *Wnt3* expression at the tip of the hypostome; salmon arrows to ectopic *Wnt3* expression in tentacles, along the body column, at basal extremity. Schematic views of *Hydra* polyps on the right depict the typical expression profile of each condition. s.t.: short tentacles; scale bars: 200 µm. **Supplement to Figure 3B, 3C.**

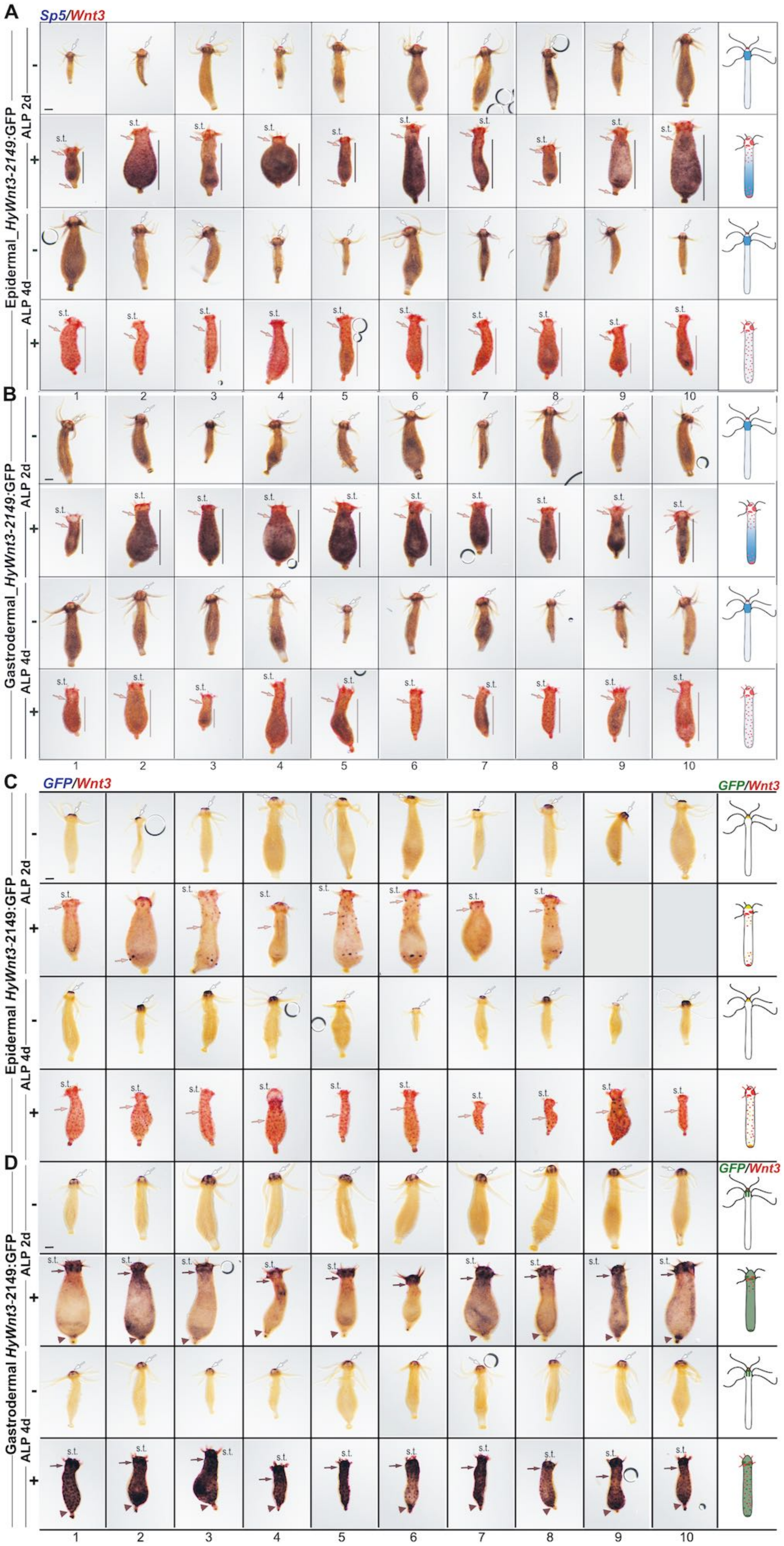


**Figure S8. Impact of ALP treatment on *GFP, Wnt3* and *Sp5* expression pattern in epidermal and gastrodermal *HyWnt3-2149*:GFP transgenic animals.**

**(A, B)** Co-detection of *Sp5* (dark blue) and *Wnt3* (red) in epidermal (A) or gastrodermal (B) *HyWnt3-2149:*GFP animals treated with DMSO (-) or ALP (+) for 2 or 4 days. In both panels, vertical bars indicate *Sp5* expression along the body column; white arrows, *Wnt3* expression at the tip of the hypostome; salmon arrows: ectopic *Wnt3* expression in the tentacles, along the body column or at the basal extremity.

**(C, D)** Co-detection of *GFP* (dark blue) and *Wnt3* (red) in epidermal (C) or gastrodermal (D) *HyWnt3-2149:*GFP animals treated with DMSO (-) or ALP (+) for 2 or 4 days. In both panels, white arrows indicate *GFP* expression at the tip of the hypostome; brown arrows *GFP* expression at the base of the apical region; salmon arrows ectopic *Wnt3* expression in the body column and brown triangles *GFP* expression at the basal extremity. For each row, a schematic view of the expression profile typical of the experimental condition is shown on the right; s.t.: short tentacles; scale bars: 200 µm. **Supplement to Figure 3B, 3C.**

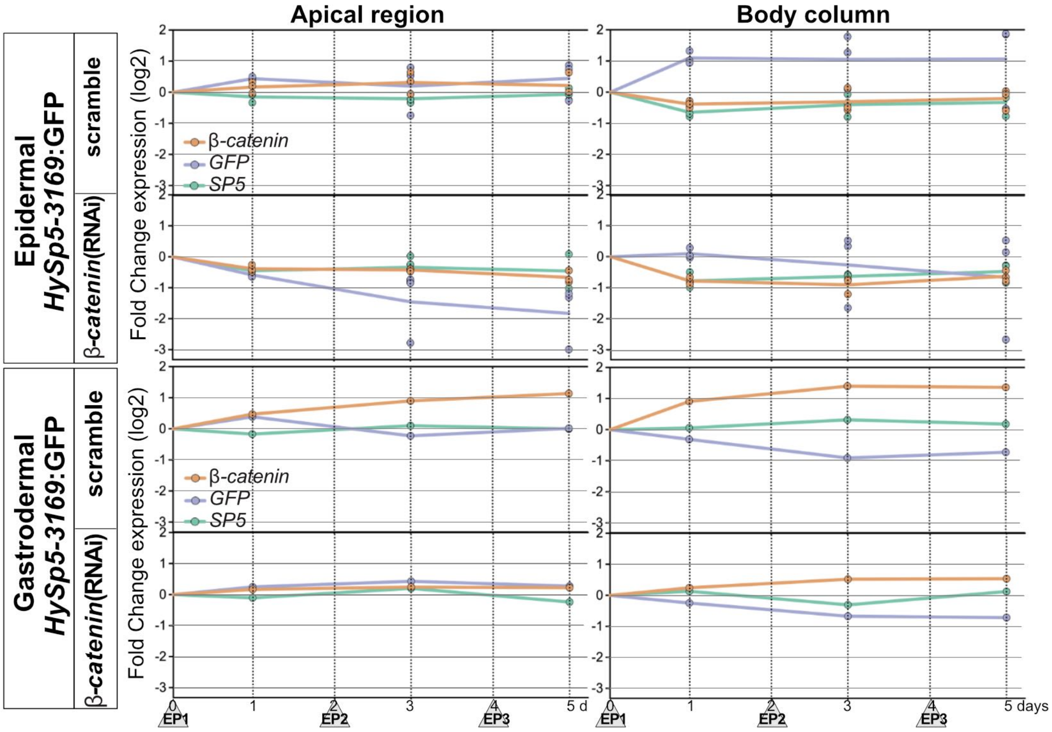


**Figure S9. Impact of *β-catenin*(RNAi) on *β-catenin*, *Sp5* and *GFP* transcript levels in epidermal and gastrodermal *HySp5-3169*:GFP transgenic animals**

Quantitative-PCR (qPCR) quantification of *β-catenin*, *GFP* and *Sp5* transcripts measured one day after EP1, one day after EP2, or one day after EP3 in *HySp5*-3169:GFP animals exposed to scramble or *β-catenin* siRNAs. Values are expressed as Fold Change (log2) when compared to the value measured in non-electroporated *Hv_AEP2* animals taken at time 0, just before (RNAi) animals are submitted to EP1. Three independent experiments were performed on epidermal *HySp5-3169:GFP* transgenic animals and one experiment on gastrodermal *HySp5-3169:GFP* transgenic animals. Supplement to Figure 4B. See raw values in supplemental Excel file.

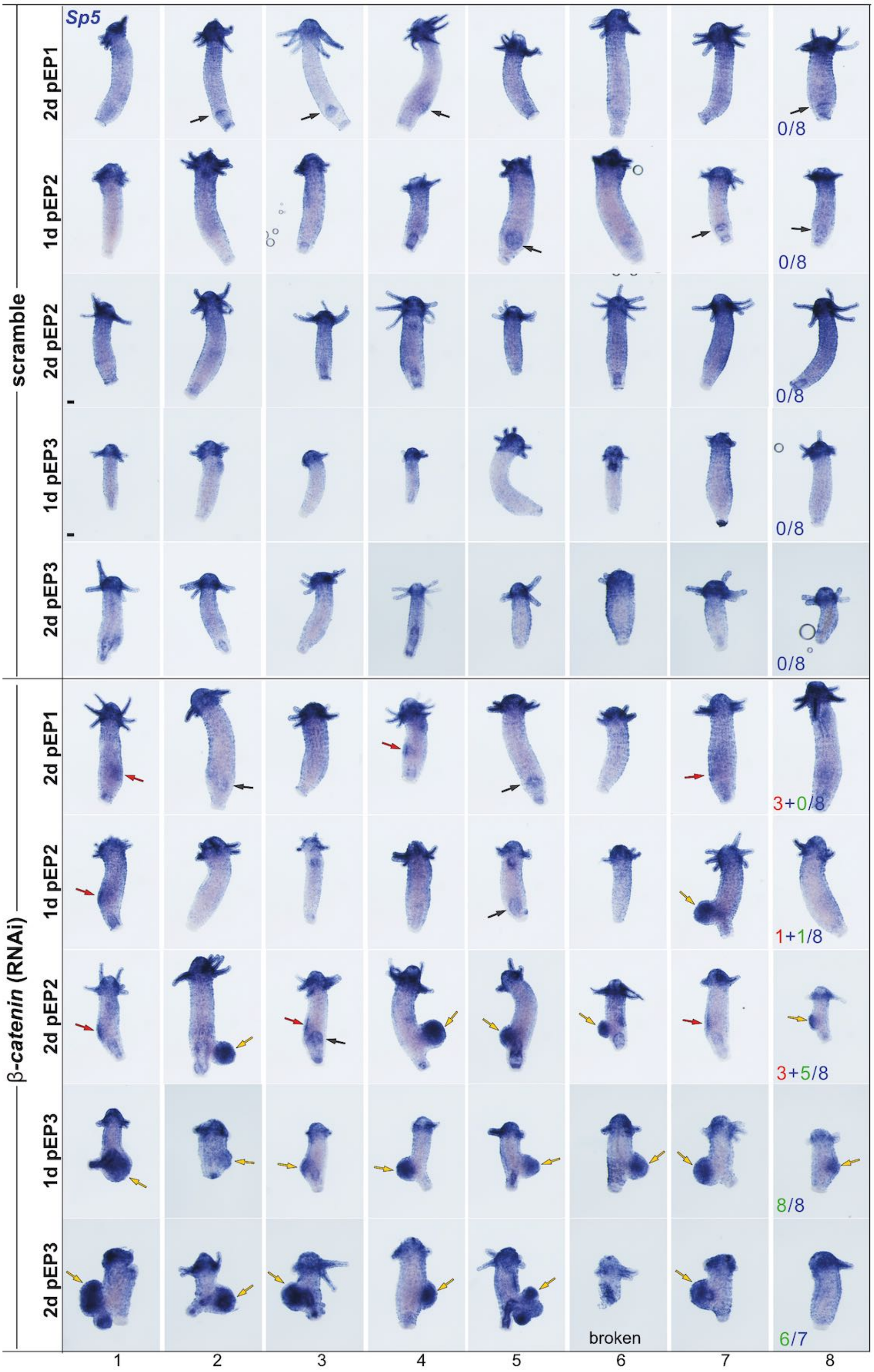


**Figure S10. Impact of *β-catenin*(RNAi) on pseudo-bud formation and *Sp5* expression in *Hv-Basel* animals**
*Sp5* expression in *Hv_Basel* animals electroporated one to three times (EP1, EP2, EP3) with scramble or *β-catenin* siRNAs and fixed as indicated: 2 days post-EP1 (2dpEP1), 1d or 2d post-EP2, 1d or 2d post-EP3. Black arrows point to detached buds, red arrows to *Sp5*-expressing patches along the body column, yellow arrows to *Sp5*-expressing pseudo-bud structures, which become multiple at 2 days post-EP3 (2dpEP3). Scale bars: 200 µm. **Supplement to Figure 4C.**

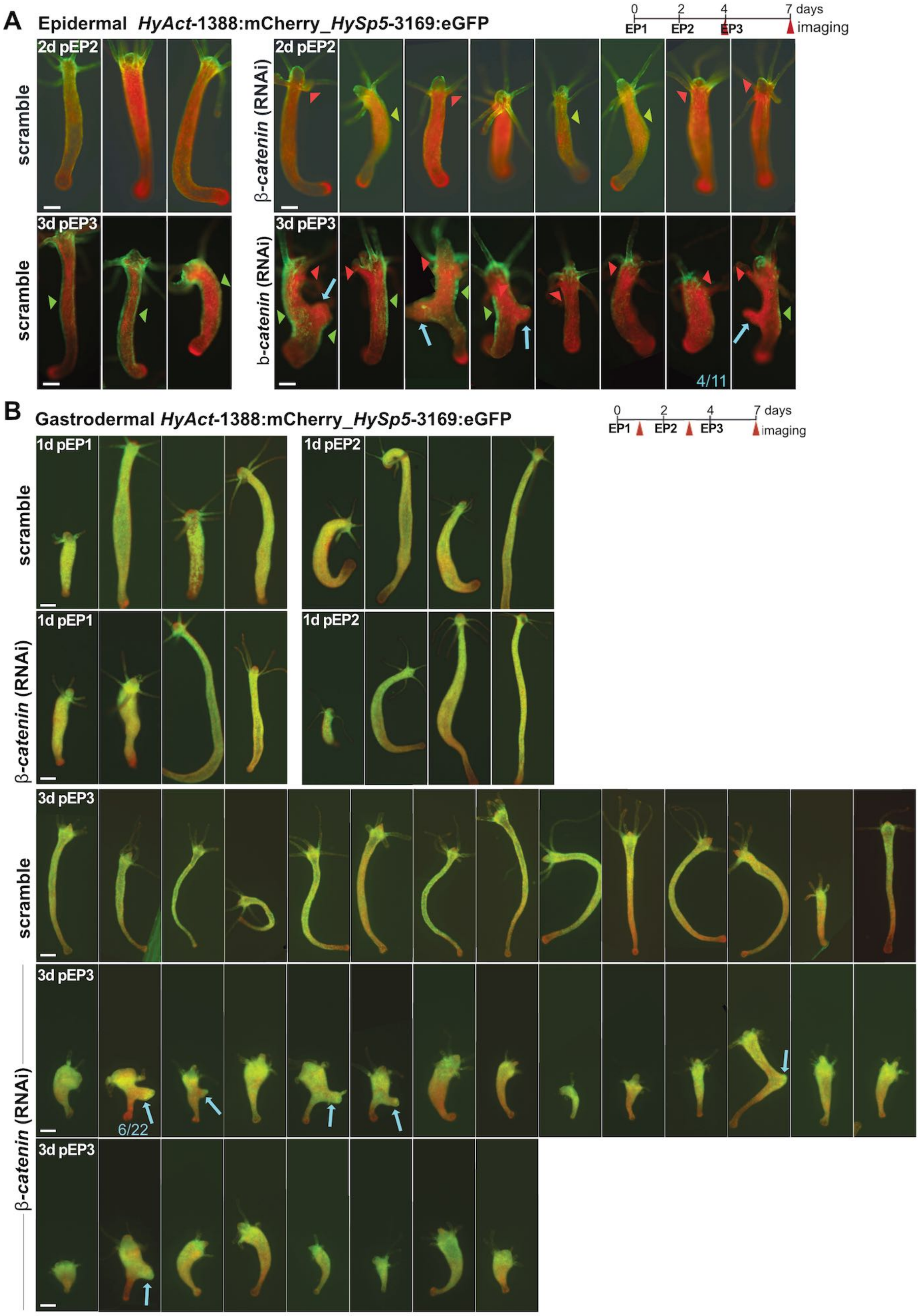


**Figure S11. Impact of *β-catenin*(RNAi) on GFP and mCherry fluorescence in epidermal and gastrodermal *HySp5-3169*:GFP transgenic animals**

GFP (green) and mCherry (red) fluorescence in epidermal **(A)** and gastrodermal **(B)** *HySp5*-3169:GFP transgenic animals electroporated with scramble or *β-catenin* siRNAs and pictured live as indicated. Red triangles indicate apical areas where GFP fluorescence is reduced, green arrowheads and green bars areas of ectopic GFP fluorescence along the body column, and blue arrows the pseudo-buds that develop at mid-body in *β-catenin* (RNAi) animals, 4/11, in epidermal *HySp5-3169*:GFP animals and all GFP-negative animals (A), 6/22 in gastrodermal *HySp5-3169*:GFP animals and all GFP-positive (B). Supplement to Figure 4C.

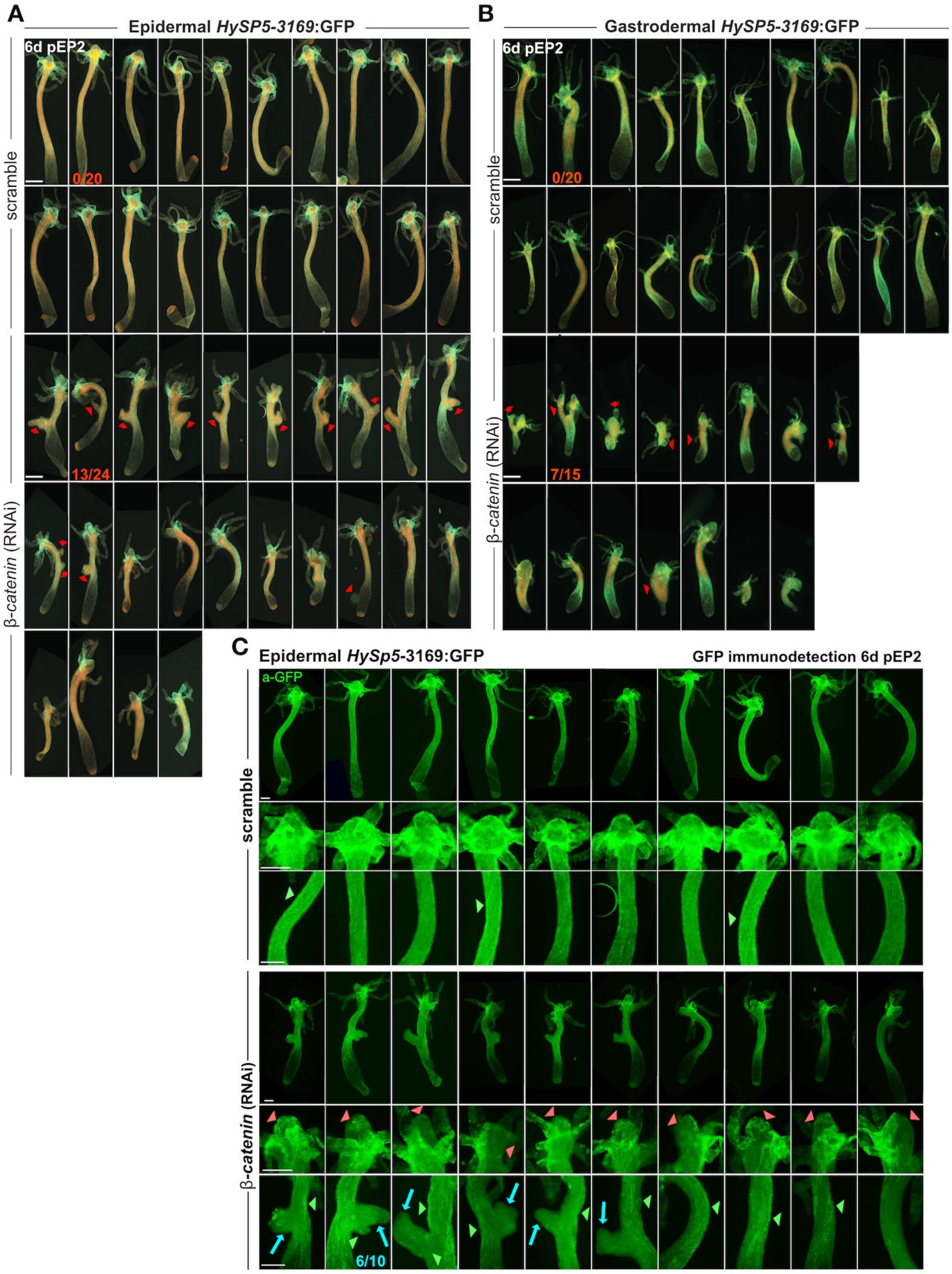


**Figure S12. Impact of *β-catenin*(RNAi) on GFP and mCherry immuno-detected patterns in epidermal and gastrodermal *HySp5-3169*:GFP transgenic animals**

**(A, B)** GFP (green) and mCherry (red) immunodetected in epidermal_ (A) and gastrodermal_ (B) *HySp5*-3169:GFP transgenic animals six days after the second electroporation (6d pEP2) with scramble or *β-catenin* siRNAs. Red arrowheads indicate pseudo-buds that develop along the body column. **(C)** GFP immunodetected in epidermal_*HySp5*-3169:GFP transgenic animals 6 days after the 2$^{nd}$ electroporation (6d pEP2) to scramble or *β-catenin* siRNAs. Enlarged views of the apical region and body column are shown for each condition. Note that in *β-catenin* (RNAi) animals GFP is largely reduced in apical regions (red triangles), absent from the pseudo-buds (blue arrows) and present in some limited areas along the body column (green triangles). Scale bars: 250 µm. Supplement to Figure 4D.

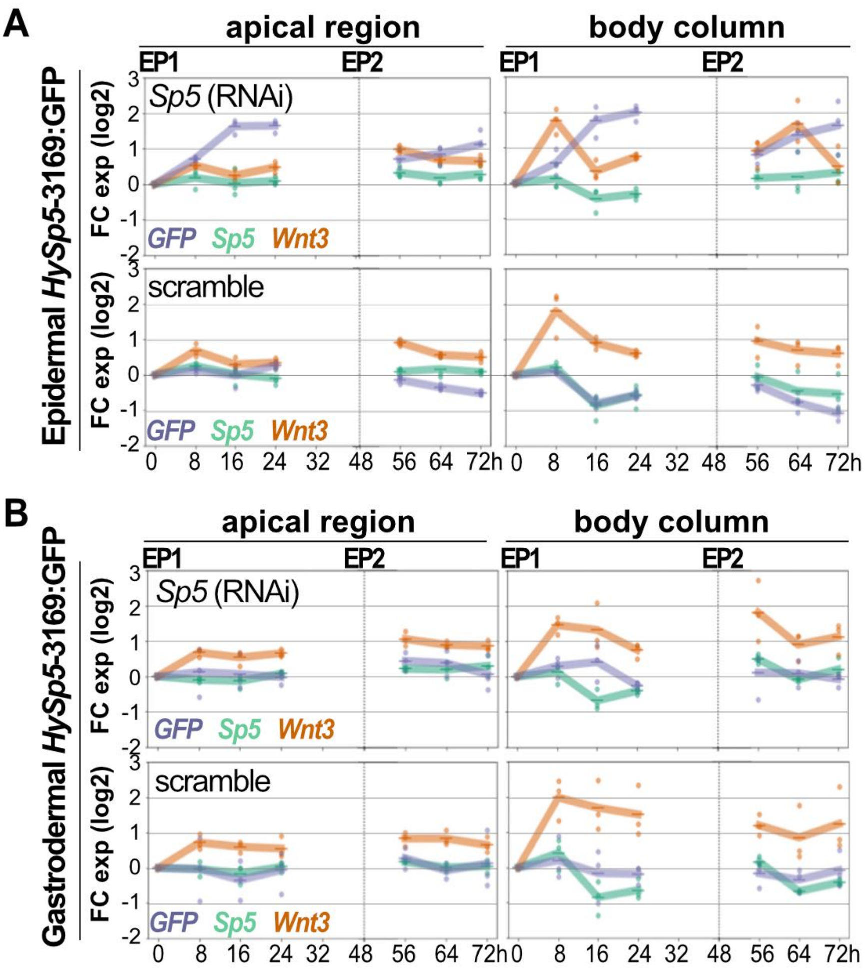


**Figure S13. Impact of *Sp5*(RNAi) on *Sp5, GFP* and *Wnt3* expression levels in epidermal and gastrodermal *HySp5-3169*:GFP transgenic animals**

Quantitative PCR analysis of *Sp5, Wnt3* and *GFP* expression levels in the apical (100%-80%, left) and body column (80% - 0%, right) regions of epidermal **(A)** or gastrodermal **(B)** *HyAct1388*:mCherry_*HySp5-3169:*GFP transgenic animals. These animals were electroporated once or twice (EP1, EP2) with scramble or *Sp5* siRNAs and imaged 8, 16, 24 hours post-EP1 and post-EP2 as described in Fig. 5A. In both panels, the colored lines correspond to the Fold Change values (FC log2) obtained by dividing values measured in *Sp5*(RNAi) or scramble animals by the reference value obtained in non-electroporated *Hv_AEP2* animals (reference at time 0, just before EP1). Note the marked increase in *GFP* expression in the apical region and body column of *Sp5*(RNAi) epidermal *HySp5-3169:*GFP transgenic animals. See raw values in supplemental Excel file. **Supplement to Figure 5A**.

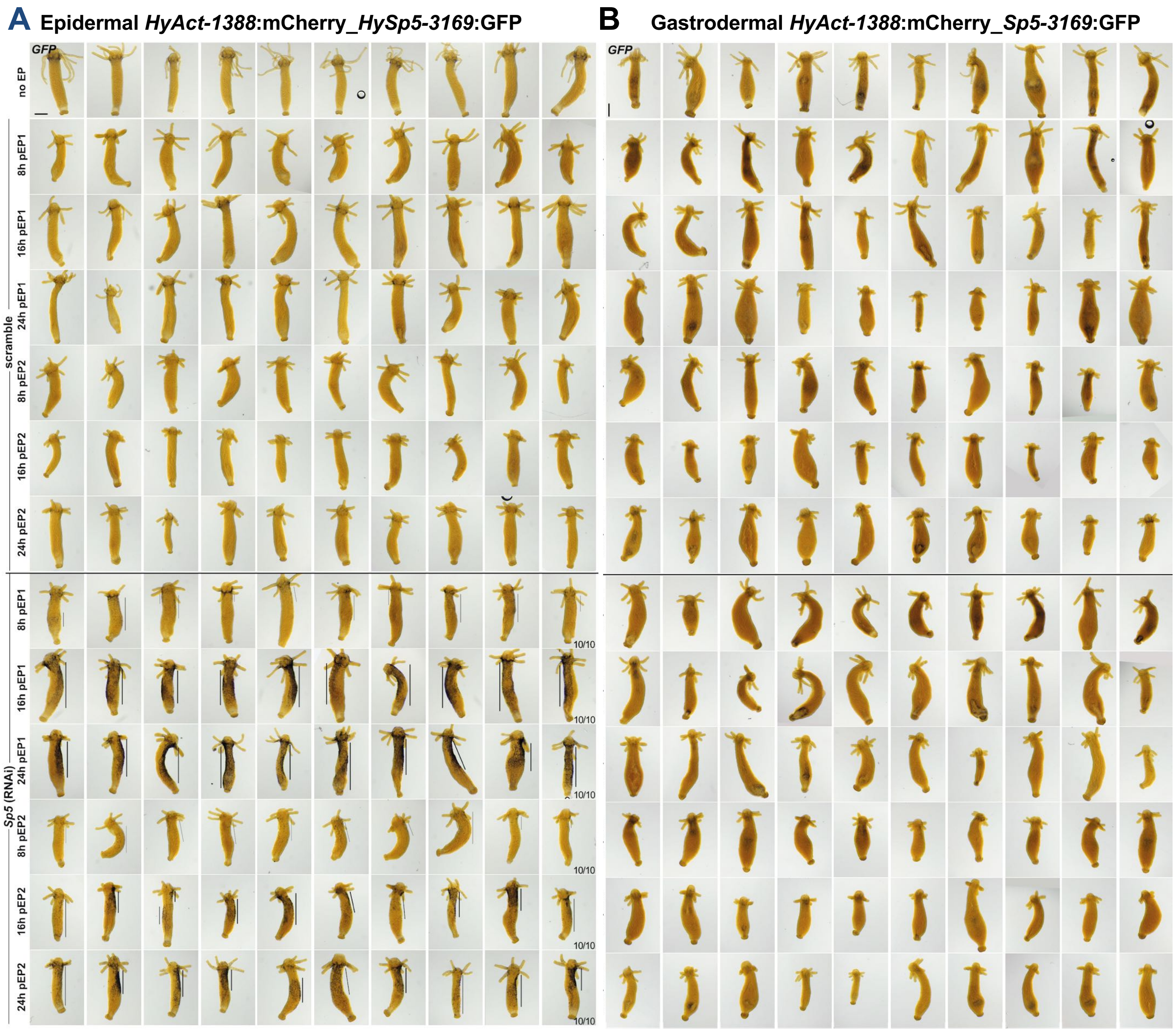


**Figure S14. Impact of *Sp5*(RNAi) on *GFP* expression in epidermal and gastrodermal *HySp5-3169*:GFP transgenic animals**

*GFP* expression in *HySp5-3169*:GFP animals either left untreated (wt, topmost row), or electroporated once or twice (EP1, EP2) with scramble or *Sp5* siRNAs and fixed for WM-ISH at 8, 16, 24 hours post-EP1 (pEP1) and 8, 16, 24 hours post-EP2 (pEP2) as described in Figure 5A. **(A)** In epidermal *HySp5-3169*:GFP animals, note the ectopic *GFP* expression along the body column (vertical bars). **(B)** In gastrodermal *HySp5-3169*:eGFP animals, no significant modulations can be observed in *Sp5* (RNAi) conditions. Scale bars: 250 µm. **Supplement to Figure 5B.**

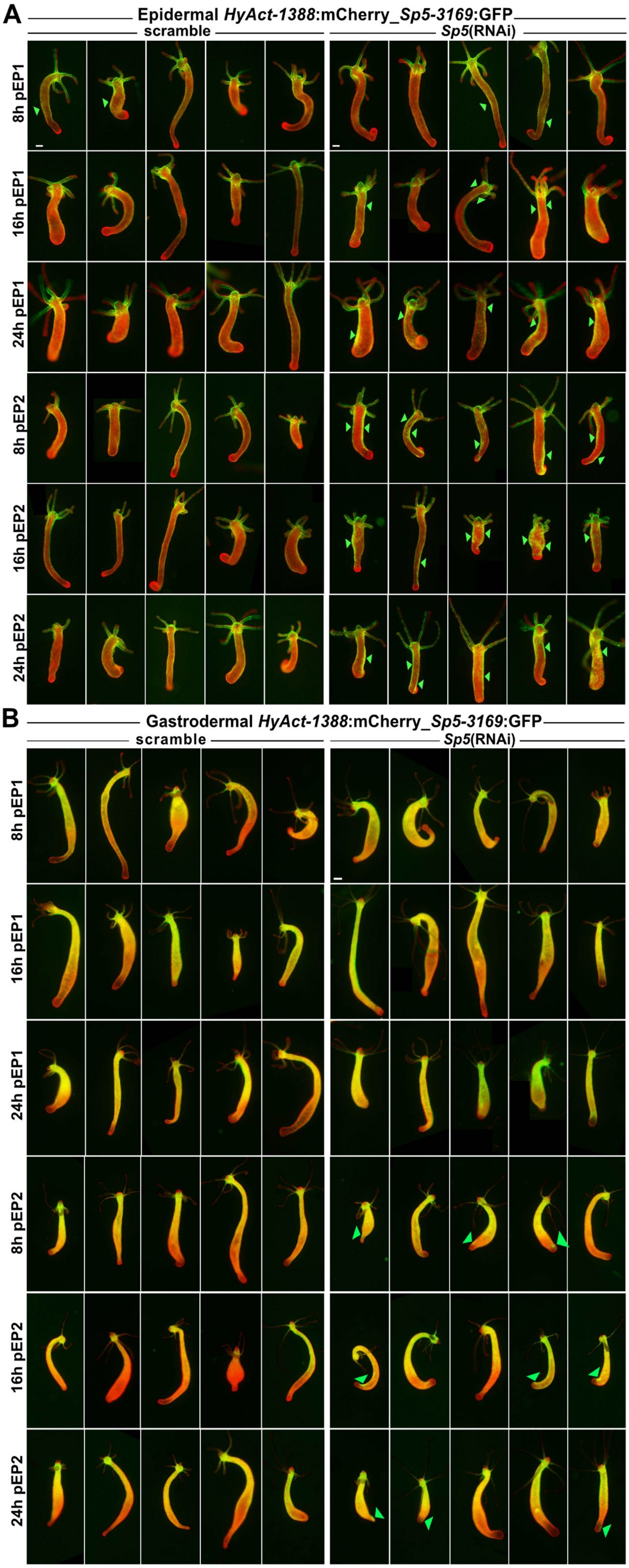


**Figure S15. Impact of *Sp5 (RNAi*) on GFP and mCherry fluorescence in epidermal or gastrodermal *HySp5-3169*:GFP transgenic animals**

Animals were electroporated once (EP1) or twice (EP2) with scramble or *Sp5* siRNAs and imaged 8, 16, 24 hours post-EP1 (pEP1) and 8, 16, 24 hours post-EP2 (pEP2) as described in Figure 5A. Images from GFP (green) and mCherry (red) merged channels are shown. Green triangles indicate areas of ectopic GFP fluorescence along the body column. Scale bars: 200 µm. **Supplement to Figure 5C.**

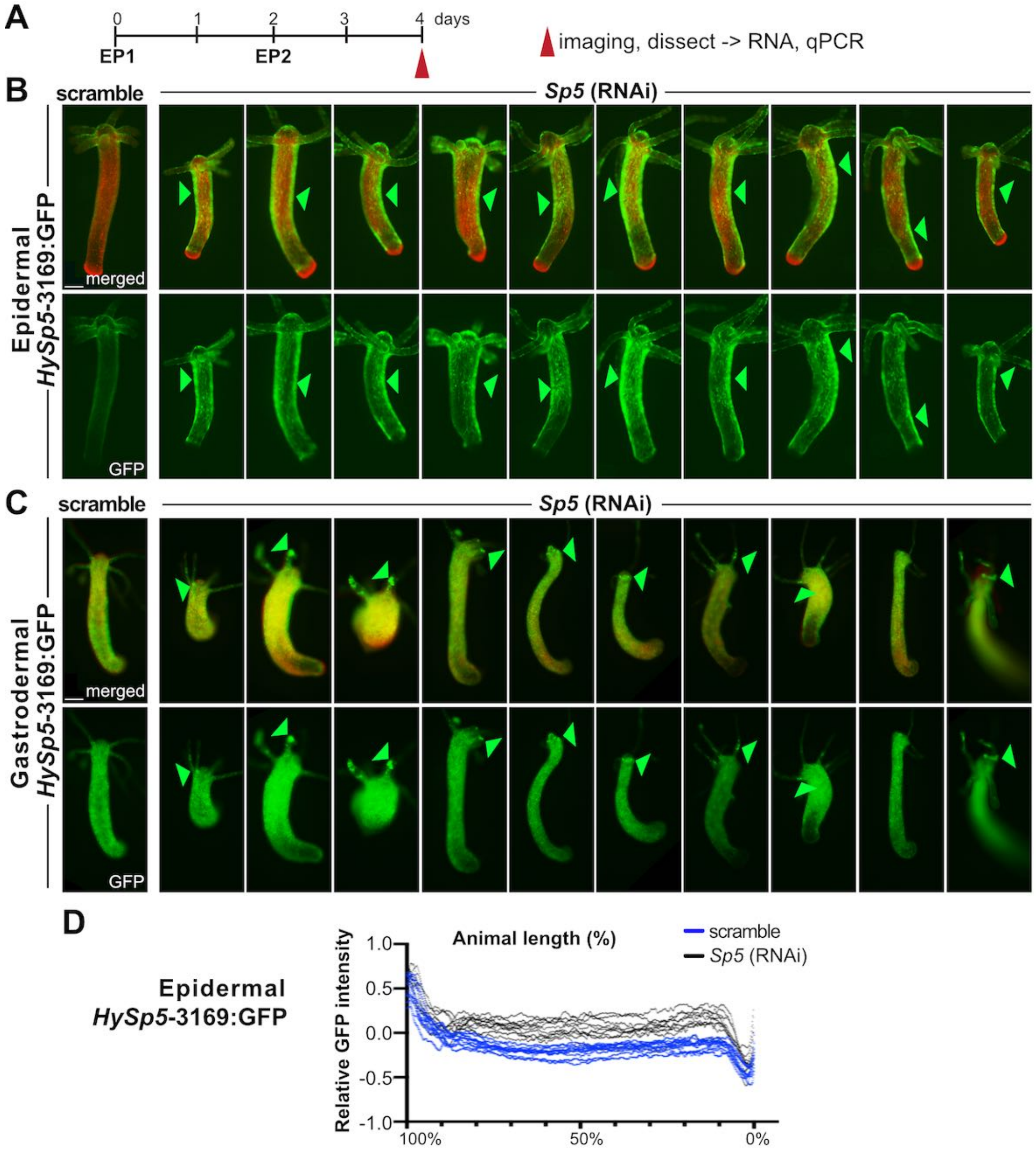


**Figure S16. Impact of *Sp5(RNAi)* on GFP fluorescence in epidermal and gastrodermal *HySp5-3169*:GFP animals**
**(A)** Schematic view of the procedure where animals are exposed twice to scramble or *Sp5* siRNAs and pictured two days later (2d pEP2). **(B, C)** Live imaging of epidermal (B) and gastrodermal (C) *HySp5*-3169:GFP transgenic animals performed two days after EP2. Green triangles point to ectopic areas of GFP fluorescence, as extended areas along the epidermis, or as gastrodermal spots in the tentacles. Scale bars: 250 µm. (**D**) Measurement two days post-EP2 of the relative GFP intensity in epidermal_*HySp5*-3169:GFP animals exposed to scramble or *Sp5* siRNAs (n=10). **Supplement to Figure 5E.**

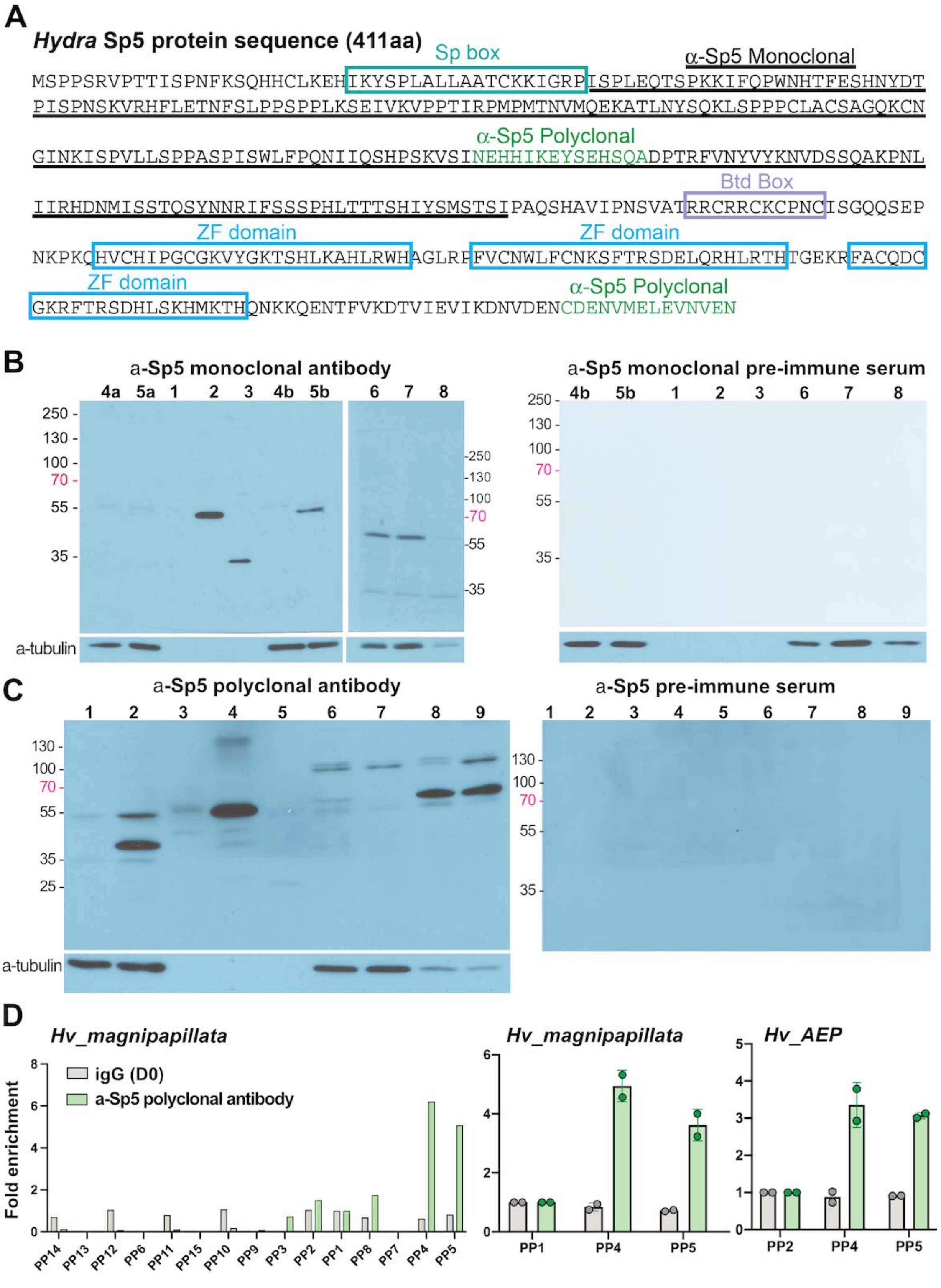


**Figure S17. ChIP-qPCR analysis of the Sp5-binding sites in the *HySp5* promoter using anti *HySp5* antibodies.** **(A)** HySp5 protein sequence with the Sp box (green), Buttonhead (Btd) box (purple) and zinc finger (ZF) domains (blue). The 218 AA long sequence of HySp5-218 protein used to raise the monoclonal anti-Sp5 antibody is underlined, the peptides used to raise the polyclonal anti-Sp5 antibody are written green. **(B)** Western blot analysis testing the anti-Sp5 monoclonal antibody (left) and the pre-immune serum (right) against the Sp5 protein either TNT-produced (lane 2; empty control: lane 1), or synthesized as recombinant HySp5-218 protein (lane 3, 24.5 kDa), or expressed in HEK293T cells (lane 5b; empty control: lane 4b), or present in *Hv_AEP2* nuclear extracts (NEs) prepared from whole animals (lane 6), apical (lane 7) or basal (lane 8) halves. **(C)** Western blot analysis testing the anti-Sp5 polyclonal antibody and the pre-immune serum against the Sp5 protein either expressed in HEK293T cells (lane 2; empty control: lane 1), or TNT-produced (lane 4; empty control: lane 3), or synthesized as recombinant HySp5-218 protein (lane 5), or present in *Hv_AEP2* NEs prepared from whole animals (lane 6), apical region (lane 7), body column (lane

8) or basal regions (lane 9). In B and C, the loading was tested by stripping the membranes and reprobing them with the anti-alpha tubulin antibody. **(D)** Enrichment in Sp5-binding along the *HySp5* promoter measured by ChIP-qPCR with *Hm-105* extracts using the anti-Sp5 polyclonal antibody and 15 primer pairs as depicted in **Figure 6C** (left), or restricted to the PP1, PP4 and PP5 regions (middle). ChIP-qPCR analysis of the PP2, PP4 and PP5 region when using *Hv_AEP* extracts and the anti-Sp5 polyclonal antibody (right). Note that in all conditions an enrichment is only detected in the PP4 and PP5 proximal regions. **Supplement to Figure 6.**

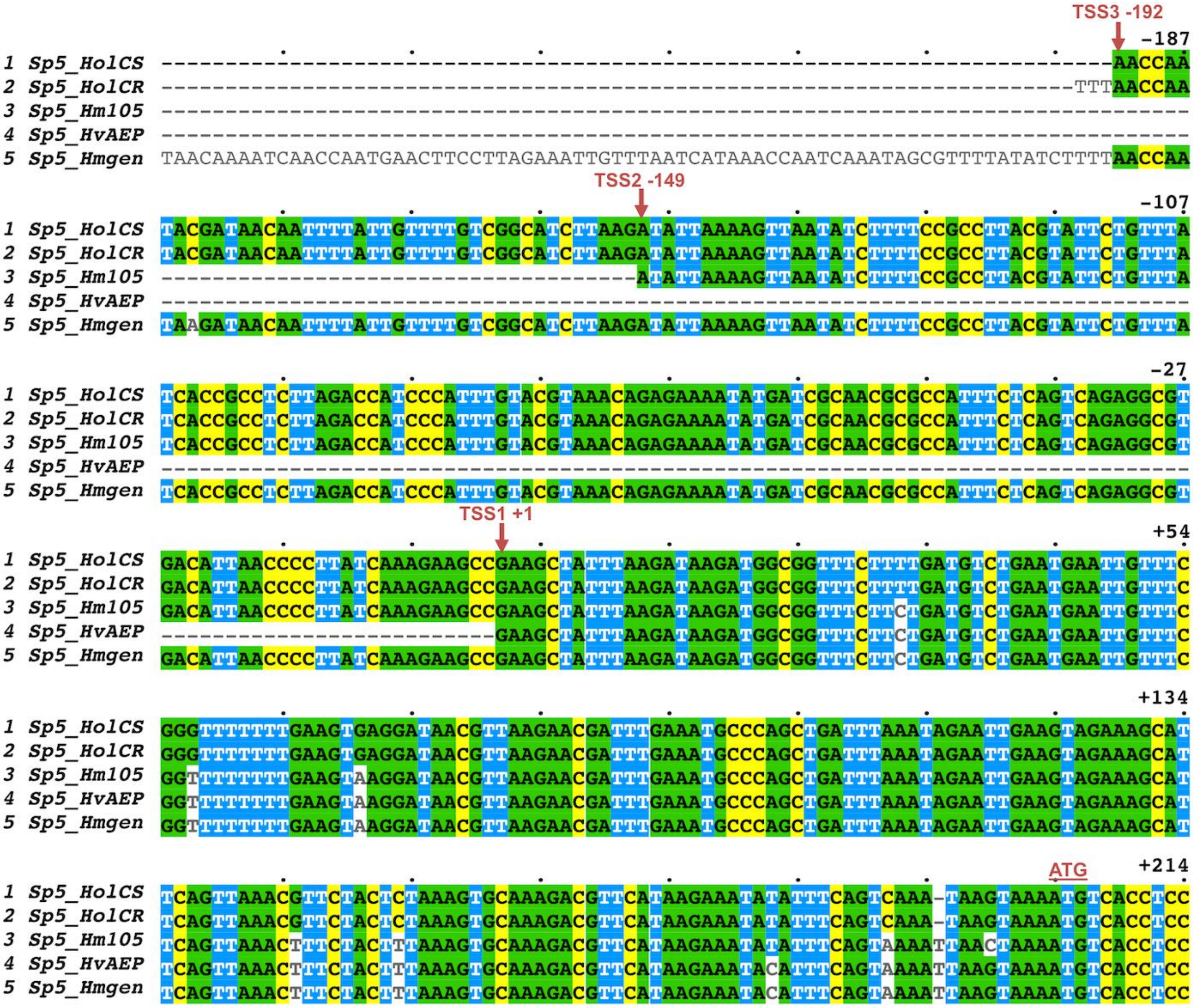


**Figure S18. Mapping the putative *Sp5* Transcriptional Start Sites (TSS)**

The *Sp5* cDNAs sequences were retrieved on HydrAtlas, https://hydratlas.unige.ch, (Wenger et al., 2019) for the Cold Sensitive or Cold Resistant *H. oligactis* (*HolCS:* S034511c0g1_i01*, HolCR*) and *H. vulgaris* (*Hv_AEP:* c16537_g1_i01) strains, on NCBI for *H. magnipapillata (Hm105:* XM_004206770.3*)*, then aligned to the *Sp5 H. magnipapillata* genomic sequence (Sp5_Hmgen) with Muscle Align (www.ebi.ac.uk/Tools/msa/muscle) selecting a ClustalW output format and finally visualized with MView 1.63 (www.ebi.ac.uk/Tools/msa/mview) [9]. Putative transcriptional start sites (TSS, indicated with vertical red arrows), were deduced from the 5' end of cDNAs from *Hv_AEP* (TSS1, position +1), *Hm105* (TSS2, position -149) and *H. oligactis* (TSS3, position -192). The *Sp5* cDNA sequence from *Hv_Jussy* (seq62049_loc 21222) was not included in the alignment as too divergent. **Supplement to Figure 6E.**

### *(A) Hydra Wnt3 upstream sequences*

```
-2149  gagttgaagagcaagaaaacgaatgattgtaaattgtacatgtcgattgtgaaaattgactataaataattattaaaaat
-2069  gtagattttatgtgcgaggaaaacatgaagaccatagaaaactccaaagcactgatgttcactgaaaaaactagataaat
-1989  aaaacgattttaactaagaggaacagtaaaaagatgcaacttagctaaatgatgagtaacgcgaaatagagttttggttg
-1909  atggaaaaaatatcgatcgctaacttgaacagcgaacatgtagtttttgttgctactggataccaaaccatggtaataat
-1829  gctatcctttttttaattttaattttgactattttttaaatcttaagcccgcgctctccatcggtaaaaaagaagaca
-1749  atggcaaccctaaattaatgtatgtatgtaaaaggtcatcatatattaacatattataataaaatattatttatgtttta
-1669  atactatttcttatatttttgattttcataaaaaatgatttgtttttatagactactattattatcatattattttta
-1589  tatttcatttttatgtttttattactaatataactattattaatctaaaacagtggaataaacatttagaatattgttga
-1509  cttcatattcgcctaaaaagcatatgtttacatttttttattaagtaattatctttgtttggcatttaataatattaaaaa
-1429  gtattcgtcaccaaaactctgctttttcaaatttataaatttatgttaacgttaaaacaatgcatcaatgcattgaacgt
-1349  ttttttaatggaaaaattaacaattttgctatctttataagtttgtttagtttcaagttttatataatgtagcttgagtg
-1269  taaaaatattatttgagtctaacaataaatctaaaataactttttgttgatgctagattttatacatctttaacagctat
-1189  ggcgccaaaagagacagcgggtgttttattgagctacgcagttattgttttttaaattcagtctttatcaaatgttaac
-1109  aaataaaaatcctgattgcatcccaatcagctcatttggaatatgttatgaccaaccaaattgttggcaaagtagttgta
-1029  aattatgatgtcatcttcttataggaaaatgacatcagtaaataatcgtcagaaaaaaacgtcagaaaacaagtgacgta
- 949  ttataatttgtttataatatacatatatatatataaaactcatcgatgacaaaacaatgaccaataaaatattcctttg
- 869  atagagtctcctaatatgttatcttggtaatttgtacaattattttattaaatgtttatcaattaatgataaatttaaac
- 789  ttatagagatatttgcaatgttgataaaaaagatgtcgataacatattaacgaaatagtatatgaatataaacatataat
- 709  attatcatatatattatgcacatcaagttttatgtttactgacgtttccctgccttctaactcctacctgcacaaaaagc
- 629  ttgaactttatatttttaattagtataagcaaatagacgtcttaaagttaaatcgatacttctttaaaatctttacaa
- 549  acattaccatgtgaaatatttggcttaatatatcagaacagctatatcaatttcactgtaattttatcgcactgtgctta
- 469  ttattagatattcaataacccattgtgaaaaggcttaaacccgtttttactttgttgctttaaaggtttgatttaactaa
- 389  ccgctaatcattataaaaagttataaatttataaaaatttatctctgcaaaaaaagatttaacattaacacaactactac
- 309  aaaaatctatctcaggatactgtcaaaaagttaaacaggtcattaaagttaacttttgtcaaagcggtatccctattcca
- 229  aaagtttcaacgtaaccgtgtcaattctttacatagcaaatagaaagggtcaccgattaaagaaaataataaaaggattc
- 149  acacgtgctaatcgtgcgctttaaagatgactgattgtttctaactttatttgaaatttcacaagaaaagttttcacatt
-  69  aatttttaaacacaatgcaaaaataaacgcaaagaatatgcagttttaaaacatgcatttttaaagaaa
```

### *(B) Hydra Zic4 upstream and coding sequences*

```
-3270   ccatgtcggttgtgtcttatagcgacttcgactttttttatttctccaaaaaaacttccgacataaactagactttgagtaaagtcgaag
-3180   agttgtttatttatattcttctttctgttttatttgctggtaaagtttttcaataaaaattatcaactgaggattaactgcaataaaaa
-3090   agtactatctgtaaataaaataataataataataaataaaaaagatggtaaaaatttacttaagtaataataaaaaactttttttgttcat
-3000   taaaatatatatatatagttgttttgtaataattttaaattttttacaatacatatcccttatctctgttaagccaaatgttttccgtt
-2910   acgaaatccttatatattgtgttatttttaaagtttacgcgacaggaagctagtaatttaatatcttttaagtttttagattttcaagt
-2820   caatatttctcttttactaataattttatgaaaataatttgttgtttttgttttgtcgttgagattggtacaataaaacatgaaaaaggc
-2730   aaatttccgattaacaaagctttggattttgataaagaaaaaaaaaatccaaatttattatttcacatgaaaggtgcgaaagattagcgg
-2640   agttttgttttgtttttgaaataatacaattacaaattctttttagaaaatgcggattagaatttattaaaatgacattaacactgtttt
-2550   gagtctaagaaacgtaaaatatgctagttatttttacaacaagcaggtaaaacaataaaaatggcctttaaagtaattttgagtttattt
-2460   cattgttgttagctcaatcgatactagttcagtaaaatacgtttatttaaagtattttactaaaattatgctctttaatattcgaaaaaa
-2370   gaaaacaaactttttccaaaaatctaacatttttcattactcaaaaaaaaactttctttttgcgctttacgcagcgctacaaaaaatatg
-2280   tcgctaagtactaaaattaaaaaaaacaaaaaaacaacgtaattacataacatgacgtaatgcaactttaatacgcatttgcagtgaata
-2190   ctttattatttatatatatatatatatactcacacatacgtatacatatatatgtataatcaaatacgtaaatagatatagatataatat
-2100   atgaatatattataagtttaatttattctattgttattgattttaaaagaaagtcagaaaatcttaaactgaaagcaaaaaaaaaactt
-2010   tttcaataatatactttttatcattattttgttgttgttgttatgattgttattataattataataataactttttattattgttattatta
-1920   ttattattgtaatcgtttatgaaaacttgatggttgctaaagatttaaggtaaataaatcgccccctctcgctattgatttgtttatttt
-1830   aaataacgggcattggcgtgaaattaatggcagcttatcatacgaattgctatacattcaaagaattttttgttagtacccatgagtttat
-1740   tttgaggtgacccatgctgttctcattaatctaaaaccattggtttttttgattaaaaggatcatcgctcaaataaagtgttttactcagg
-1650   cgtgtgataactgtattactaagacttcctcgatataaacttgcatcaaattatttctgaataatgattattgtcaaaaacaaagacaca
-1560   gttgactaaattcctaacttactttgtattatttttttgtctgtaattttatagaccgatatgttactttttgctacatgtgttagaagttg
-1470   tttgagaattcaaactgagtgaaaaaaatggacaaataaaacaaaataaaacgcgtatgtcctgctaccttttttatagctgttttaaata
-1380   aagtttcttttggtattctaatgttgtgacttctaaaaaaattatgcaatcaaagattatcgacgcaacgataagtcaattaaaggaaag
-1290   gggtgctacatatgagacaacacatgtgtgttttacattttttaaccggtttaacgtcactccgaacaaagtttaaattaatgcgaacgtt
-1200   ttatttttgtagctaaacatctgagttaaaaatgcttgcgcacgttcttttaacaaaaacagctcatcttattgttttactttttattttt
-1110   ttattaaattaaaaaaaatataattaaaaaatattttgtaagaatcttaacagatatttacgataaaatttcttatctaaagtatcgt
-1020   tttcaggttactcgcgtagacttataattttcaataaagattacccaatttgtctaattagcataacaattaattataagtaaataaccg
- 930   gagataataagtaaataacctgagataaaaacctcaaagagtacagcagattggaaatcatttttttttttttcaagaaaccgcgcgcat
- 840   tcactcgtataccttaattgtataatcttttagttaaacaagcaccaaaaccggaacaaaggtagtagtatacaatcaaagcaagctttg
- 750   atatctctaatattttttttcaattttttttttttatccacttatttgtctgagctatttctttaccttaattacgctataaaatttttcttct
- 660   cgacaagcggcgtagctttgagtttatagctcttttctgaagtctaattccttctgtttttattacaatcctgactaagcttcctcaaag
- 570   tcaaggaaaacaaatgcagttaaaaagaaaataatccttgtaaacacgtttcatccttcataaaaaacgtacgctaataaagctttttttg
- 480   tttacagttaagcaagctaatttagaatttcagaattactctttgcaaaaaaaaaaaaaaaaaaaaaaaaaaaaaaaaaaaaaaaaa
- 390   agtaataataataattagccacgtcattttgcaaccagattaaaatccaggaaatctcaatgaaattatttgggaggaagttgtatattt
- 300   ctaaaatttagcagaacttatattagcgtatatcttcgacacacataaacttactgtgatattacaaaatgactcaattgagttagaaat
- 210   aaaatcaaataaataaactaaatttcccaatactacaaagcaggatctgaaaggttcttttaaggattctcgcttcatacttttttttgt
- 120   tttgtattatttttgctcccgttaattaggaaataaactaccatgcagattaattatgggtataattgaaagttcgtgcagttttattctt
                                               (Hv_Jussy) TSS1
-  30   cgtcaaattaaatcaataatttagttctatttcgttttattactataaatcctctaaagaacattttttgcttgttttctttttaaagatc
Hv_Ju                                        TTCGTTTTATTACTATAAATCCTCTAAAGAACATTTTTGCTTGTTTTCTTTTTAAAGATC
                                                                      (Hol_CS) TSS2
+  60   tcattttaattttaataccccgctatgatgtgattggatgggatcaaatgtgattggttaaaaaagtttgaataagtatcaagaactgt
Hv_Jus  TCATTTTAATTTTTAATACCCCGCTATGATGTGATTGGATGGGATCAAATGTGATTGGTTAAAAAAGTTTGAATAAGTATCAAGAACTGT
Hol_CS  ------------------------------------------GTCGAATGTGATTGGTTAAAAAAGTTTGAATAAGTATCAAGAACTGT
```

*(Hol_CR, Hv_AEP1)* **TSS3**

+ 150 caatcaaaataaatgagcggccaatcaggttagttataaatccttattagcaacataatccctacacatagacagatgtaccatacataa
*Hv_Jus* CAATCAAAATAAATGAGCGGCCAATCAGGTTAGTTATAAATCCTTATTAGCAACATAATCCCTACACATAGACAGATGTACCATACATAA
*Hv_AEP* -------------------------------------------------------------------------------------**CA**TAA
*Hol_CS* CAATCAAAATAAATGAGCGGCCAATCAGGTTAGTTATAAATCCTTATTAGCAACATAATCCCTACACATAGACAGATGTACCATACATAA

+ 240 tggaaaggtaaaattgatttgtgcccaaaagcatttaaaagtctcaatttacatccgaatgacaccgtatttcagtcacagtcagttgga
*Hv_Jus* TGGAAAGGTAAAATTGATTTGTGCCCAAAAGCATTTAAAAGTCTCAATTTACATCCGAATGACACCGTATTTCAGTCACAGTCAGTTGGA
*Hv_AEP* TGGAAAGGTAAAATTGATTTGTGCCCAAAAGCATTTAAAAGTCTCAATTTACATCCGAATGACACCGTATTTCAGTCACAGTCAGTTGGA
*Hol_CS* TGGAAAGGTAAAATTGATTTGTGCCCAAAAGCATTTAAAAGTCTCAATTTACATCCGAATGACACCGTATTTCAGTCACAGTCAGTT**A**GA

+ 330 agcagcgttctagacgtatcagagggttatctagttgataattgaatgtgacattgttttattaaatataaaagtatatttttatgcagc
*Hv_Jus* AGCAGCGTTCTAGACGTATCAGAGGGTTATCTAGTTGATAATTGAATGTGACATTGTTTTATTAAATATAAAAGTATATTTTTATGCAGC
*Hv_AEP* AGCAGCGTTCTAGACGTATCAGAGGGTTATCTAGTTGATAATTGAATGTGACATTGTTTTATTAAATATAAAAGTATATTTTTATGCAGC
*Hol_CS* AGCAGCGTTCTAGACGTATCAGAGGGTTATCTAGTTGATAATT**A**AATGT**A**ACATTGT**G**TTATTAAATATAA**C**A**T**TAT**T**TT**C**TTATGCA**A**C

+ 420 gtttgtattgtggtgtt-------acctcattaaatatctacaaattgttcctatattaaaacctaattatttatctagaaacttattta
*Hv_Jus* GTTTGTATTGTGGTGTT-------ACCTCATTAAATATCTACAAATTGTTCCTATATTAAAACCTAATTATTTATCTAGAAACTTATTTA
*Hv_AEP* GTTTGTATTGTCGTGTT-------ACCTCATTAA**G**TATCTACAAATTGTTCCTATATTAAAACCTAATTATTTATCTAGAAACTTATTTA
*Hol_CS* **A**TTTGTAT**A**GTGGTGTTA**AA**TCATACCTC**G**TTAAATATCTACAAATT**A**TTCCT**T**TATTAAAACCTA**G**TT**G**TTTAT**TGG**GAA**G**CTTATTTA

+ 510 ttaaatatctgtttcatta-aa**atg**gagcatcgtttacaaagttcaacattaggacattatccaaacgttccacggtactattacttaga
*Hv_Jus* TTAAATATCTGTTTCATTA-AAATGGAGCATCGTTTACAAAGTTCAACATTAGGACATTATCCAAACGTTCCACGGTACTATTACTTAGA
*Hv_AEP* TTAAATATCTGTTTCATTA-AAATGGAGCATCGTTTACAAAGTTCAACATTAGGACATTATCCAAACGTTCCACGGTACTATTACTTAGA
*Hol_CS* TT**T**AATATCTGTT**A**CATT**GA**A**C**ATGGAGCATCGCTTGCAAAGTTCAACATTAGGACATTATCCAAACGTTCC**G**CG**A**TATTATTACTTAGA
M E H R L Q S S T L G H Y P N V P R Y Y Y L D

### Figure S19. Putative *Zic4*-binding sites in the *Wnt3* and *Zic4* genomic sequences from *Hm-105 Hydra*

**(A)** Putative Zic-binding sites in the 2’142 bp *Wnt3* genomic sequence from *Hm-105* animals at positions -1781 and -1177 (highlighted in blue). **(B)** Putative Zic-binding sites in the 3’420 bp *Zic4* upstream genomic sequences (written lowercase) from *Hm-105* animals at positions – 1863, - 1824, -1744, -1289, -107 and +77 (highlighted in blue). This sequence is available in the Zic4-2505:GFP reporter construct positions 5’200 up to 8’993 (www.addgene.org/193001) [29]. *Zic4* cDNA sequences from two *H. vulgaris* strains (*Hv_Jussy, Hv_AEP1*) or *H. oligactis* (CS strain) are written uppercase. See accession numbers in **Table S1**.
